# Structure of the Xylan *O*-Acetyltransferase AtXOAT1 Reveals Molecular Insight into Polysaccharide Acetylation in Plants

**DOI:** 10.1101/2020.01.16.909127

**Authors:** Vladimir V. Lunin, Hsin-Tzu Wang, Vivek S. Bharadwaj, Markus Alahuhta, Maria J. Peña, Jeong-Yeh Yang, Stephanie A. Archer-Hartmann, Parastoo Azadi, Michael E. Himmel, Kelley W. Moremen, William S. York, Yannick J. Bomble, Breeanna R. Urbanowicz

**Author notes:** Corresponding Authors: Yannick J. Bomble and Breeanna R. Urbanowicz. These authors contributed equally to this work (Vladimir V. Lunin, Hsin-Tzu Wang, Vivek S. Bharadwaj, Markus Alahuhta).

## Abstract

Acetylation of biomolecules is gaining increased attention due to both the abundance and importance of this modification across all kingdoms of life. Xylans are a major component of plant cell walls and are the third most abundant biopolymer in Nature. *O*-Acetyl moieties are the dominant backbone substituents of glucuronoxylan in dicots and play a major role in the polymer-polymer interactions that are crucial for proper wall architecture and normal plant development. Here, we describe the biochemical, structural, and mechanistic characterization of Arabidopsis thaliana xylan O-acetyltransferase 1 (AtXOAT1), a member of the plant-specific Trichome Birefrigence Like (TBL) family that catalyzes the 2-*O*-acetylation of xylan. A multipronged approach involving X-ray crystallography, biochemical analyses, mutagenesis, and molecular simulations show that XOAT1 catalyzes xylan acetylation through formation of an acyl-enzyme intermediate by a double displacement bi-bi mechanism involving a Ser-His-Asp catalytic triad and unconventionally employs an arginine residue in formation of an oxyanion hole.

## Introduction

The plant cell wall provides a structural scaffold that surrounds each plant cell, defining the size and shape of plant cells and tissues. The plant cell wall is a composite material comprised of cellulose microfibrils, hemicellulosic polysaccharides, pectins, proteins, glycoproteins, and aromatic molecules (including lignins). Xylans, which constitute a major portion of hemicellulose, are one of the most abundant plant polysaccharides on earth and are hence considered to be key targets for biomass modification to enhance the production of biofuels and bioproducts (Smith et al. 2017). Xylans, like most cell wall polysaccharides, with the exception of mixed-linkage glucan and cellulose, are *O*-acetylated (Pauly and Ramírez 2018). All xylans produced by vascular plants are decorated with glycosyl, or acyl groups on an identical backbone structure composed of 1,4-linked β-D-xylopyranosyl (Xyl) residues. The nature and pattern of decorations on xylan vary depending on the plant tissue and species (Smith et al. 2017). For example, in dicots, *O*-acetylated glucuronoxylan (AcGX) is the most abundant hemicellulosic polysaccharide and consists of a 1,4-linked β-D-xylopyranosyl backbone that is substituted with 1,2-linked α-D-glucuronic acid (GlcA) and/or its 4-*O*-methyl derivative (MeGlcA). Approximately 60% of the backbone residues are substituted with acetyl groups, making them the most abundant substituents of xylan. More detailed analysis of xylan acetylation in Arabidopsis (Col-0) showed that 44% of the xylosyl residues are monoacetylated at *O*-2 or *O*-3, 6% are bisubstituted at both *O*-2 and *O*-3, and approximately 75% of the (Me)GlcA substituted backbone residues are also acetylated at the *O*-3 position (Chong et al. 2014). Further analysis showed that acetyl residues are present on every alternate Xyl along the backbone where glycosyl substituents are absent, and (Me)GlcA residues are localized according this alternating acetylation pattern (Chong et al. 2014), suggesting that acetylation may play a key role in the systematic addition of substituents in a defined pattern along the polymer backbone.

While the specific structural and biological roles of polysaccharide acetylation in the plant cell wall remain enigmatic, *O*-acetylation is thought to be a key player in determining the hydrophobicity of AcGX, and hence a key determinant in the interactions between xylan and other wall polymers, including cellulose and lignin (Johnson et al. 2017; Busse-Wicher et al. 2014). For example, the presence of O-acetyl substituents decreases adsorption of xylan to cellulosic surfaces, indicating that the interactions between xylan- and cellulose and xylan and - lignin interactions could be modulated by the degree and patterning of substituent decorations(Köhnke et al. 2011; Kang et al. 2019). Recent studies on *Populus* genotypes with different cell wall compositions suggest that there is a close interaction between lignin and xylan, and that the degree of xylan acetylation influences the interaction between these major cell wall polymers, thus impacting pretreatment efficiencies (Johnson et al. 2017). Furthermore, acetyl groups sterically hinder hydrolytic enzymes, and thus decrease the accessibility of those enzymes to their polysaccharide targets (Biely 2012). Taken together, it is evident that acetylation of xylans affects cell wall architecture and mechanical strength through its interaction with other polymers in the cell wall. Therefore, insights into acetylation mechanisms in plants form the basis for a deeper understanding into the interplay between polysaccharide functionalization and cell wall architecture, growth and development. Advances in this area may be used to overcome biomass recalcitrance to enzymatic saccharification, leading to the development of improved design schema for engineering efforts or targeted genomics approaches for the conversion of cell wall rich plant biomass into sustainable bioproducts.

Until recently, little was known about the process of plant cell wall polysaccharide *O*-acetylation. Despite the importance of this modification in both cell wall structure and organization, the identity of the acetyl donor substrates or the existence of donor intermediates and the exact roles of many of the proteins involved in polysaccharide acetylation remain unknown; however, much progress is being made in this area. So far, it has been shown that at least four protein families are involved in the acetylation pathway in plants, including the Trichome Birefringence-Like (TBL) protein family, Reduced Wall Acetylation (RWA) proteins (Manabe et al. 2013), the Altered XYloglucan 9 (AXY9) protein (Schultink et al. 2015), and a newly described GDSL acetylesterase family (Pauly and Ramírez 2018) that will not be discussed herein.

Members of the TBL family have been shown to function as polysaccharide *O*-acetyltransferases and participate in the *O*-acetylation of specific cell wall polymers (Yuan et al. 2015; Yuan et al. 2016b; Zhong et al. 2017, 2018; Stranne et al. 2018). ESKIMO1/TBL29 (At3g55990) is one of the most well characterized members of this family. Mutation of this gene in Arabidopsis (*eskimo1, tbl29-1, tbl29-2*) results in plants with collapsed xylem vessels (Lefebvre et al. 2011), that are tolerant to salt, drought and freezing stresses. Analysis of the cell walls of these mutants showed that they produce xylan with 50-60% less *O*-acetylation, indicating that this enzyme is a key player in xylan acetylation (Xiong et al. 2013). Our earlier studies on this enzyme provided direct biochemical evidence showing that TBL29/ESK1 is an xylan specific *O*-acetyltransferase that catalyzes the addition of *O*-acetyl groups to the 2-position of xylosyl backbone residues *in vitro*, establishing the precise molecular function of this enzyme(Urbanowicz et al. 2014), and lead to the proposed name xylan *O*-acetyltransferase 1 (XOAT1). Recently, an eloquent report by Grantham and coworkers showed that the even pattern of xylan acetylation is absent in the *esk1* mutant, indicating that AtXOAT1 is necessary for patterning of acetyl esters on xylan in Arabidopsis (Grantham et al. 2017). Since the initial biochemical analysis of AtXOAT1, several other members of the TBL family have also been shown to play a role in the regiospecific acetylation of xylan in Arabidopsis, including TBL3 (XOAT4), TBL28 (XOAT2), TBL30 (XOAT3), TBL31 (XOAT5), TBL32 (XOAT6), TBL33 (XOAT7), TBL34 (XOAT8), and TBL35 (XOAT9) (Yuan et al. 2016b; Yuan et al. 2015; Yuan et al. 2016a; Zhong et al. 2017).

In plants, acetyl-CoA cannot readily cross membranes and is independently produced in plastids, mitochondria, peroxisomes and the cytosol, but not in the Golgi apparatus. Members of the RWA family function non-specifically in polysaccharide *O*-acetylation at a biosynthetic step prior to TBLs and have been proposed to transport an activated form of acetate into the Golgi using cytosolic acetyl-CoA pools (Manabe et al. 2013). However, none of the multi-transmembrane proteins or domains of known polysaccharide *O*-acetylating systems, including RWA, OatA, and AlgI have been shown to function as acetyl-CoA transporters that transport acetyl-CoA across a membrane (recently reviewed by (Brott et al. 2019; Pauly and Ramírez 2018)). These proteins are currently proposed to function by translocating acetyl groups derived from cytosolic acetyl-CoA or another yet unknown donor in both plants and microbes, but not acetyl-CoA itself, across membranes(Brott et al. 2019). AXY9 has been suggested to play a role in the polysaccharide *O*-acetylation process subsequent to the action of RWAs but prior to that of TBLs. Similar to the RWAs, AXY9 functions non-specifically in polysaccharide *O*-acetylation and is thought to play a role in the production of an intermediate acetyl donor substrate. Interestingly, AXY9 contains the GDS and DxxH motifs found in members of the TBL family (Schultink et al. 2015). Recently, AXY9 was shown to possess weak acetylesterase activity using the psudosubstrates 4-methylumbelliferyl acetate (4MU-Ac) and *p*-nitrophenyl acetate(Zhong et al. 2018). Taken together, these data suggest that AXY9 may be able to use acetyl donors other than acetyl-CoA to form an acyl- -AXY9 intermediate, which may serve as an proteinaceous activated acetyl donor itself or as an intermediate step in the formation of an as yet to be determined acetyl donor.

Understanding the precise function of enzymes involved in plant polysaccharide biosynthesis, including glycosyltransferases, acetyltransferases, and methyltransferase at the molecular level is essential for gaining fundamental insight into how these biocatalysts work together to build architecturally complex structures such as the cell wall. Several key studies have laid the foundation regarding the biochemical function of XOAT1 as an *O*-acetyltransferase involved in xylan biosynthesis, indicating its key role in substituent patterning, thought to be critical for the wall architecture. Structural information of enzymes involved in polysaccharide biosynthesis, especially those involved in the addition of methyl and acetyl substituents present in plant cell wall polysaccharides, are completely absent in structural databases despite their significant roles in plant growth and development. Traditionally, this was due to difficulties producing sufficient quantities of these plant-derived enzymes for crystallization trials. Recently, mammalian cell expression has been successfully used to express plant glycosyltransferases involved in xyloglucan biosynthesis in sufficient quantities for structural and biochemical characterization, including *Arabidopsis thaliana* fucosyltransferase-1 (*At*FUT1) and xyloglucan xylosyltransferase (*At*XXT) (Urbanowicz et al. 2017; Culbertson et al. 2018). Herein, we applied this strategy for structural characterization of the xylan *O*-acetyltransferase *At*XOAT1. We showed that the structure of *At*XOAT1 is characterized by a deep cleft on the surface of the protein separating the molecule into two unequal lobes and a Ser216-His465-Asp462 catalytic triad is located at the bottom of this cleft. Molecular simulations of AtXOAT1 in its substrate bound states and experimental characterization of the acyl-enzyme intermediate provide evidence for a double-displacement acetylation mechanism.

## Results

### Biochemical insights into *At*XOAT1 catalysis

Members of the plant-specific domain of unknown function (DUF) 231 family, also referred to as the trichome birefringence-like (TBL) family are Golgi-localized polysaccharide *O*-acetyltransferases (Pauly and Ramírez 2018). *At*XOAT1 is comprised of a single NH_2_-terminal transmembrane (T) domain followed by an NH_2_-terminal variable region (NV), an N-terminal Cys-rich PMR5 domain (PF14416) that is part of the plant-specific TBL region, the DUF231 domain, and contains six predicted *N*-glycosylation sites (Fig. 1a). Previously, we developed a robust mass spectrometry (MS)-based xylan *O*-acetyltransferase assay (Fig. 1b) using acetyl-CoA as the acetyl donor and 2-aminobenzamide β-1,4-xylohexaose (Xyl_6_-2AB) to study enzymes involved in polysaccharide *O*-acetylation *in vitro (18)*. The peptidoglycan *O*-acetyltransferase from *Neisseria gonorrhoeae* PatB and the secondary cell wall polysaccharide *O*-acetyltransferase from *Bacillus cereus* PatB1 were recently shown to display donor substrate promiscuity and utilize chromogenic acetyl-donor substrates *in vitro(Brott et al. 2019; Sychantha et al. 2018)*. To determine whether other activated acetyl substrates may function as donor substrates for *At*XOAT1, we performed *in vitro* xylan acetylation assays to compare the ability of *p*-nitrophenyl acetate (*p*NP-Ac), acetylsalicylic acid (ASA), acetyl-CoA and 4-methylumbelliferylacetate (4MU-Ac) to function as donor substrates for *At*XOAT1. Matrix-assisted laser desorption ionization time-of-flight mass spectrometry (MALDI-TOF MS) analysis of the reaction products indicated that recombinant *At*XOAT1 catalyzes the transfer of *O*-acetyl moieties from all four substrates to Xyl_6_-2AB. Compared to acetyl-CoA, *p*NP-Ac, ASA and 4MU-Ac all serve as better donor substrates *in vitro* (Fig. 1c). This is consistent with a recent report by Ye and colleagues that was published during the preparation of this manuscript (Zhong et al. 2019).

**Figure 1.**
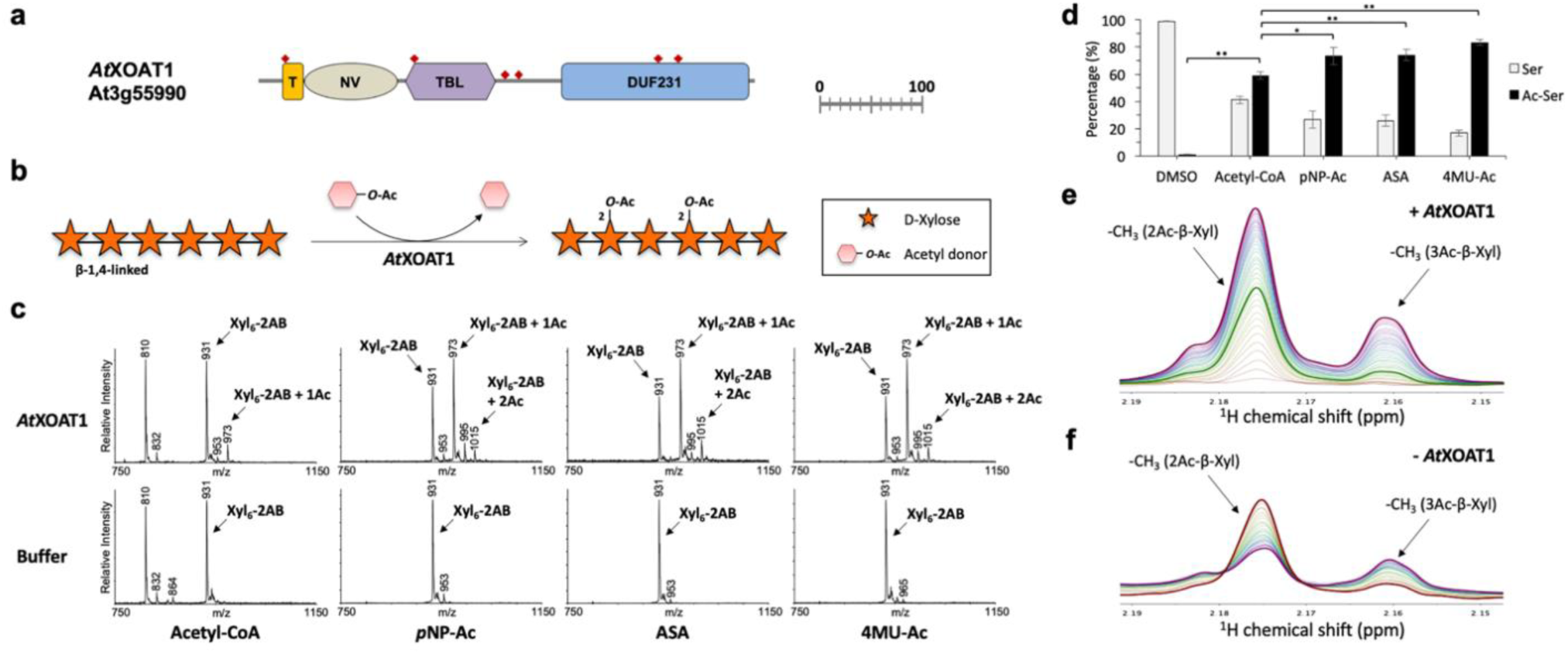
Biochemical properties of *At*XOAT1. (**A**) The scaled domain architecture of *At*XOAT1 described in this study. T, NH_2_-terminal transmembrane domain; NV, NH_2_-terminal variable region; TBL, TBL domain; DUF231, DUF231 domain. The transmembrane helix was predicted using TMHMM Server v. 2.0 (http://www.cbs.dtu.dk/services/TMHMM/; Krogh et al., 2001). Prediction of the *N*-glycosylation sites (red diamonds) were carried out using the NetNGlyc 1.0 Server (http://www.cbs.dtu.dk/services/NetNGlyc/; Blom et al., 2004). Scale bar represents 100 amino acids. (**B**) The *in vitro* reaction scheme of xylan *O*-acetylation catalyzed by *At*XOAT1. (**C**) The donor substrate promiscuity of *At*XOAT revealed by MALDI-TOF MS analysis of acetylated Xyl_6_-2AB generated by incubating *At*XOAT1 with Xyl_6_-2AB acceptors and different acetyl-donors, acetyl-CoA, *para-*nitrophenyl acetate (*p*NP-Ac), acetylsalicylic acid (ASA) and 4-methylumbelliferylacetate (4MU-Ac) for 6 h. Each transfer of *O*-acetyl group increases the mass of Xyl_6_-2AB by 42 Da as indicated by annotating [M+H]^+^ ions. (**D**) The percentage of acyl-*At*XOAT1 formed upon reaction with different acetyl donors. The error bars indicate mean ± s.d. from three independent assays. Statistical significance was determined with the two-tailed Student’s *t*-test. *, *p* < 0.05; **, *p* < 0.01. (**E**) Real-time ^1^H NMR analysis of *At*XOAT1-catalyzed reaction by using acetyl-CoA and xylohexaose (Xyl_6_) as the substrates. The thickened green line corresponds to the spectrum of the reaction at the 6 h time point, while the thickened purple line indicates the end of the 20-hour reaction. (**F**) Acetyl group migration analysis through real-time ^1^H NMR after removal of *At*XOAT1. The thickened red line corresponds to the starting point for monitoring acetyl group migration after removal of AtXOAT1 after a 6-hour reaction, while the thickened purple line represents the end point of monitoring the migration (14 hours after transfer).

The preference for non-acetyl-CoA co-substrates is consistent with what was observed for PatB (Moynihan and Clarke 2013), and calls into question the identity of the natural acetyl donors for this enzyme family *in planta*. However, the observation that *At*XOAT1 is able to utilize surrogate compounds as donor substrates facilitated the development of an *in vitro* xylan acetyltransferase assay utilizing the chromogenic acetyl donor *p*NP-Ac, adapted from methods described for bacterial polysaccharide acetyltransferases(Brott et al. 2019). We performed spectrophotometric assays to evaluate the *O*-acetyltransferase activity of *At*XOAT1 by using *p*NP-Ac as a donor substrate together with increasing concentrations of xylopentaose (Xyl_5_) as an acceptor to validate the robustness of our assays (Fig. S1). The release of *p*NP increased in parallel with the concentration of Xyl_5_ added to the reactions, indicating that the presence of Xyl_5_ facilitates the activity of *At*XOAT1 as a xylan *O*-acetyltransferase and confirmed that this is a robust method to determine the biochemical properties of xylan acetyltransferases. Next, we evaluated the ability of *At*XOAT1 to use xylobiose and xylooligosaccharides with degrees of polymerization (DP) from two to six as acceptors, and showed that xylotriose is the smallest acceptor that *At*XOAT1 is able to *O*-acetylate (Fig. S2), consistent with prior biochemical analyses of *At*XOAT1 using ^14^C-labeled acetyl-CoA, confirming the robustness of the spectrophotometric assay (Zhong et al. 2017).

### *At*XOAT1 *O*-acetylates xylan through the formation of a covalent acetyl-enzyme intermediate

Next, we sought to determine whether *At*XOAT1 can catalyze the formation of a covalent acetyl-enzyme intermediate with different donors in the absence of acceptor substrate. Therefore, to determine whether an acyl-enzyme intermediate is formed during the reaction, we incubated different donors with *At*XOAT1 in the absence of acceptor. The enzyme was then digested by trypsin and the fragments analyzed by LC-MS/MS. The obtained peptides containing the catalytic Ser residue S216 (210-MMFVGD**<u>S</u>**LNR-219) appeared as a mixture of the acetylated and unmodified peptides with a difference in molecular weight of 42 Da (shifted by *m/z* 2 for the doubly charged ion where *z* = 2; Fig. S3) corresponding to an attachment of an acetyl group. The sequences of the peptides were mapped by MS/MS analysis, showing that the putative active Ser residue, S216, was the acetylation site, as shown in the higher energy collisional dissociation (HCD) fragmentation spectra (Fig. S4). Further analysis of the peak areas of the extracted ion chromatographs (XIC) indicated that about 59% of the acetylated peptide population was formed when acetyl-CoA was used as a donor substrate for *At*XOAT1, while 73%, 74% and 83% of peptide populations were acetylated when *p*NP-Ac, ASA and 4MU-Ac were used as donors, respectively (Fig. 1d). It is worth noting that the preferential donor substrates for *At*XOAT1 revealed by the results of acyl-enzyme intermediate analysis are consistent with our MALDI-TOF MS analysis of the products of the *in vitro* assays (Fig. 1c), revealing that acetyl-CoA is the least preferred donor for *At*XOAT1 catalysis. Furthermore, the formation of an acetyl-enzyme intermediate at S216 suggested a double displacement mechanism for *At*XOAT1-catalyzed acetyl transfer.

### *At*XOAT1 is an obligate 2-*O*-acetyltransferase

The spontaneous migration of acetyl groups has been proven to occur on xylooligosaccharides between the *O*-2 and *O*-3 positions of xylosyl backbone residues (Kabel et al. 2003), and can even rapidly migrate to position 4 on the non-reducing end(Biely et al. 2004). This makes it difficult to examine the regiospecificity of *O*-acetyltransferases by solely analyzing the products after the reaction is completed(Zhong et al. 2017). Therefore, the interpretation of data regarding polysaccharide acetylation can be difficult, and it is unclear whether equilibrium conditions observed are present in the native plant polysaccharide itself or are established during isolation procedures. Indeed, Kabel et al. (Kabel et al. 2003) have specifically shown that freeze-drying and/or re-dissolving the isolated material promotes acetyl migration on xylooligosaccharides, confirming that the position and distribution of *O*-acetyl moieties cannot be extrapolated to the xylan structures present in the plant. Hence, the phenomenon of acetyl migration complicates determining XOAT1’s enzymatic regiospecificity. In order to unambiguously confirm the regiospecificity of these enzymes, we developed an experimental approach that that can rapidly monitor spontaneous changes in the product during catalysis and after product formation. In these assays, we measured product formation from time zero as the enzyme is functioning in real time at 5 min intervals, confirming that the enzyme is an obligate 2-*O*-acetyltransferase. Studies by Ye et al. relied on measuring the final product after long reaction times (16 hrs), which would be highly affected by spontaneous migration (Zhong et al. 2017).

*At*XOAT1-catalyzed *O*-acetylation of xylohexaose (Xyl_6_) was monitored by real-time ^1^H NMR spectroscopy using an optimized protocol to effectibly control the pH of the reaction (Fig. 1e), and showed that the resonance of the methyl protons of the acetyl groups attached to the *O*-2 position (δ 2.176) appeared at the beginning and dominated throughout the incubation period, confirming that *At*XOAT1 catalyzes *O*-acetylation at the *O*-2 position on xylosyl residues. This is similar trend to the one reported in our previous study (Urbanowicz et al. 2014). The resonance corresponding to the 3-*O*-acetyl groups (δ 2.160) showed up after the appearance of 2-*O*-acetyl resonance, suggesting an acetyl group migration possibly occurred in xylopyranose units as the migration phenomenon of acyl groups has been wildly observed and studied (Kabel et al. 2003; Biely et al. 2004; Lassfolk et al. 2018). To confirm the acetyl group migration occurring in our reactions is driven by a non-enzymatic mechanism, after reacting for 6 hours (bold red line), *At*XOAT1 was removed using a spin filter and the intensities of resonances corresponding to 2-*O* and 3-*O*-acetyl groups were continuously monitored using real-time ^1^H NMR spectroscopy for an additional 14 hrs (Fig. 1f). In contrast to the spectra obtained during the *At*XOAT1-catalyzed reaction, a decrease of the resonance of 2-*O*-acetyl groups was observed in parallel with an increase of 3-*O*-acetyl resonance, indicating that spontaneous migration of acetyl groups from *O*-2 to *O*-3 position occurred after *At*XOT1-catalyzed addition of acetyl groups to the *O*-2 position. These results unambiguously indicated the regiospecificity of *At*XOAT1 by minimizing the confounding effects of spontaneous acetyl group migration, which follows the enzymatic reaction.

### Crystal structure reveals atomistic architecture of *At*XOAT1

Three truncated forms of *At*XOAT1 were generated as fusion proteins containing an NH_2_-terminal signal sequence followed by an 8xHis tag, an AviTag, ‘superfolder’ GFP, and the Tobacco Etch Virus (TEV) protease recognition site followed by truncated coding regions of *At*XOAT1 (Fig. S5a and b). One construct was designed with a truncation devoid of the NH_2_-terminal cytoplasmic tail and predicted transmembrane domain (a.a 44–487, *At*XOAT1-fl), a second encodes the catalytic domain and lacks the N-terminal variable region (a.a. 133-487, *At*XOAT1-cat), and the third is solely comprised of the N-terminal variable region (a.a. 44-133, *At*XOAT1-nv). Expression and secretion of GFP-*At*XOAT1-fl (∼121 mg L^-1^) and GFP-*At*XOAT1-cat (∼92 mg L^-1^) in HEK293S (GnTI-) cells yielded high levels of secreted recombinant fusion protein based on GFP fluorescence (Table S1)(Moremen et al. 2018). The activities of *At*XOAT1-fl and *At*XOAT1-cat and were virtually the same. Reactions performed with *At*XOAT1-nv did not result in the production acetylated products, as expected (Fig. S5c), and *At*XOAT1-nv was used as a control protein for comparative analysis using chromogenic substrates in later experiments. Taken together, these data indicate that the core catalytic domain is comprised of the TBL and DUF231 domains, and confirms that the N-terminal variable region does not play a direct role in catalysis.

The crystal structure of the *At*XOAT1 catalytic domain (AtXOAT1-cat) was determined by the single wavelength anomalous dispersion method using anomalous signals from sulfur atoms (S-SAD) at CuK_α_ wavelength (1.54 Å). The *At*XOAT1-cat domain monomer has approximate dimensions of 45Åx48Åx55Å and features an overall fold that is unique based on a search for related structures using PDBeFold(http://www.ebi.ac.uk/msd-srv/ssm/ssmcite.html). Overall there are three β-sheets composed of seven (β4, β5, β6, β3, β9, β12, β13), four (β7, β8, β10, β11) and two (β1 and β2) β-strands, nine α-helices including a ‘broken’ one and three α-helical turns (4-residue fragments stabilized by a single H-bond). The protein topology is presented in Fig S6. A deep cleft divides the molecule into two unequal lobes. It is worth noting that the larger lobe formed by amino acids 197-366, 403-433 and 470-486 contains almost all secondary structure elements found in the protein – two parallel/antiparallel β-sheets, comprised of seven and four β-strands, respectively, and eight large α-helices (all helices except H8). The smaller lobe includes amino acid residues 133-196, 367-402 and 434-469 and is mostly unstructured with only one short broken α-helix (H8), three α-helical turns (αT1, αT2 and αT3) and a small two-strand antiparallel β-sheet (β1 and β2). All these unstructured loops are held together by four disulfide bridges (a.a. 140/191, 162/227, 171/467, 384/463). Interestingly, there are no disulfides present in the larger, more structured lobe (Fig. 2). The deep cleft on the surface of the protein molecule is the most likely substrate binding site, with a putative classic catalytic triad (Ser216-His465-Asp462) characteristic of serine proteases (Dodson and Wlodawer 1998) at the bottom. Accordingly, it has previously been shown that mutation of these residues abolishes *At*XOAT1 activity (Zhong et al. 2017), and our structural data now confirms that these residues are part of a classic catalytic triad. The walls of the cleft are formed by two flexible loops comprised of amino acids 443-448 on one side and 273-281 on the other.

**Figure 2:**
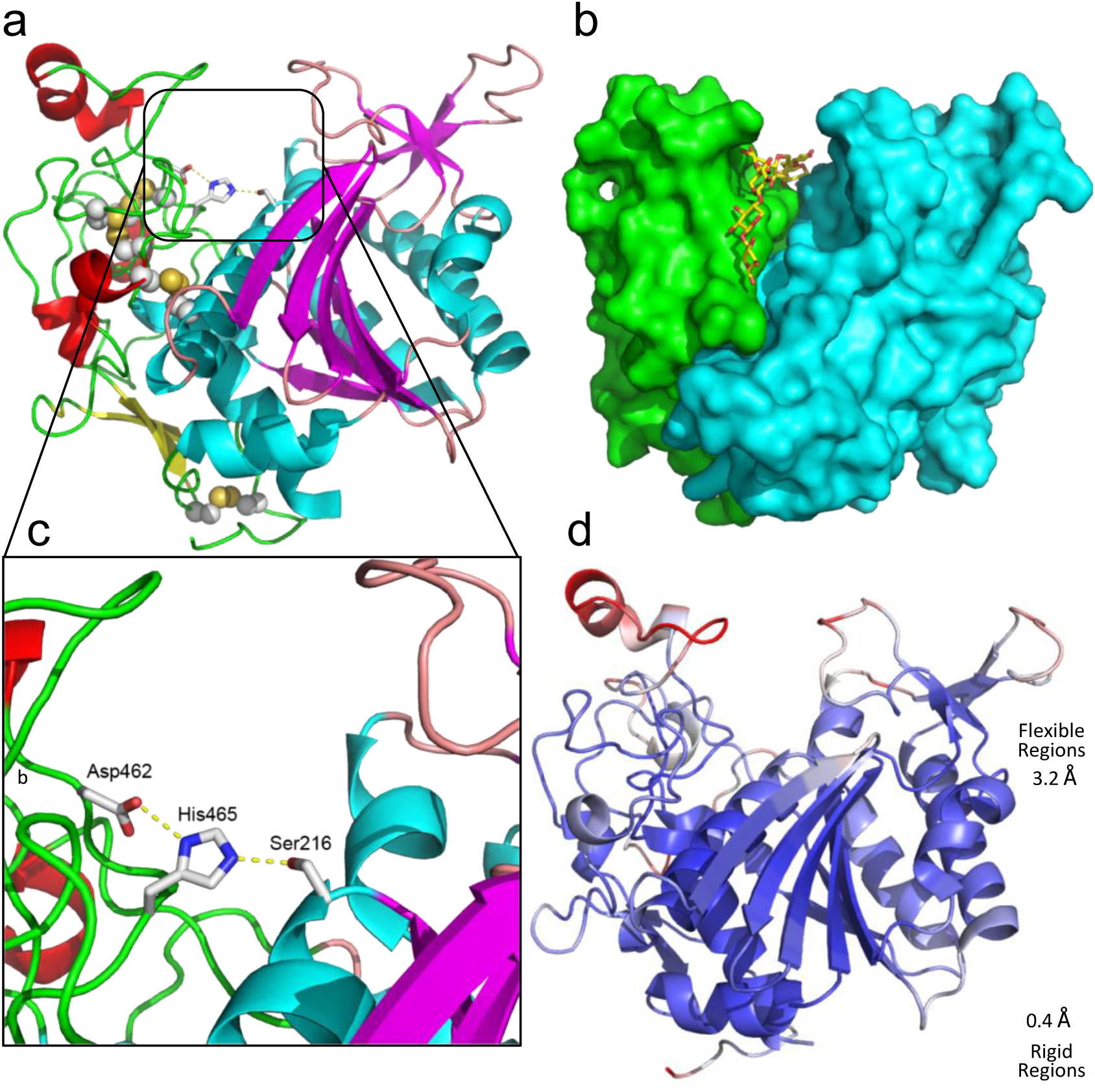
Structural topology of *At*XOAT1: (**A**) Cartoon representation of the *At*XOAT1-cat fold. Pink loops, cyan α-helices and magenta β-strands are shown for the major lobe and green loops, red α-helices and yellow β-strands are shown for the minor lobe. Disulfides are shown as spheres. The sidechains of the catalytic triad Asp462-His465-Ser216 are shown in stick representation. (**B**) Surface representation reveals a putative substrate binding groove. The major lobe is shown in cyan and the minor lobe is colored green. A xylodecaose bound in the groove in MD simulations is shown in stick representation. (**C**) Zoomed in active site. (**D**) Cartoon representation of the XOAT1-cat colored by flexibility of the protein regions based on MD simulations.

There are six *N*-glycans decorating the surface of *At*XOAT1-cat. Five of the glycans are well-defined single *N*-acetylglucosamine (GlcNAc) sugars attached to Nδ atoms of asparagine residues at positions 151, 241, 255, 393 and 412, as expected after treatment with endo-beta-N-acetylglucosaminidase F, which cleaves the β-1,4-link between the core N-acetylglucosamine residues of N-linked glycans. The sixth glycan appended to Asn425 is longer, with two GlcNAc units visible in the electron density maps that could be modeled followed by at least one mannose unit visible but not modeled due to weak and scattered electron density. Since most of the glycans are involved in the crystal contacts or are located in the immediate proximity to the crystal contacts, trimming of glycan structures to a single GlcNAc at 4 out of 5 respective sites was critical for successful crystallization of *At*XOAT1-cat.

The catalytic triad (Ser-Asp-His) located in the active site cleft of *At*XOAT1 was originally identified in serine proteases and is found in several dissimilar enzymes and protein folds that cleave amide or ester bonds via nucleophilic attack (Dodson and Wlodawer 1998). A search for similar structures using the PDBeFOLD server indicated that the overall fold of *At*XOAT1-cat is not present in any previously reported structures in the Protein Data Bank (PDB). However, some subregions of the enzyme, especially the central β-sheet region in the vicinity of the catalytic triad, have some similarities to known structures despite sharing less than 10% sequence identity. Structural alignments performed with PDBeFold found two structures with a Z-score above six – a putative lipase from *B. thetaiotamicron* (PDB ID 3bzw, Z-score 8.2, RMSD of 2.71 Å over 155 Cα atoms) and peptidoglycan *O*-acetylesterase from *N. meningitidis* (PDB ID 4k3u, Z-score 6.9, RMSD of 2.84 Å over 176 Cα atoms). Both proteins belong to the Pfam GDSL-like lipase/acylhydrolase family and share the α/β/α arrangement of *At*XOAT1-cat’s larger ‘structured’ lobe including five of the seven β-strands in the largest β-sheet flanked with seven α-helices on both sides (Fig. 3). The central β-sheet region of *At*XOAT1-cat and some of the surrounding α-helices also share some similarities upon structural alignment with two functionally similar enzymes - isoamyl acetate hydrolyzing esterase (PDB ID:3mil)(Ma et al. 2011) and peptidoglycan *O*-acetyltransferase from *B. cerus* (PATB1) (PDB ID: 5v8e)(Sychantha et al. 2018). Isoamyl acetate hydrolyzing esterase performs the hydrolysis of acetyl esters, whereas PatB1 transfers an acetyl moiety from acetylated donor molecules, such as *p*NP-Ac onto the C3 hydroxyl of terminal β-GlcNAc residues of peptidoglycan and hence is somewhat related by function to *At*XOAT1. Catalytic activity in these four similar proteins is also carried out by a conserved Ser-His-Asp catalytic triad. It is interesting to note that the order of appearance of the secondary structure domains; as well as the catalytic triad in the amino acid sequence is Ser-Asp-His (wherein Ser is closer to the N-terminus and the His is closer to the C-terminus) for the putative lipase, the peptidoglycan *O*-acetyltransferase from *N. meningitidis*, isoamyl acetate hydrolyzing esterase and *At*XOAT1. However, even though PatB1 is an acetyltransferase acting on a glycosyl moiety, the arrangement of the secondary structural elements as well as the catalytic triad is shifted to Asp-His-Ser (wherein Asp is closer to the N-terminus while the Ser is closer to the C-terminus). Hence, this structural similarity of PatB1 with *At*XOAT1 and the other similar proteins may be evidence of convergent evolution for use of a common catalytic triad, despite low primary sequence similarity among the enzymes (≤10%). This has previously been found for serine proteases containing a similar conserved catalytic triad (Buller and Townsend 2013).

**Figure 3.**
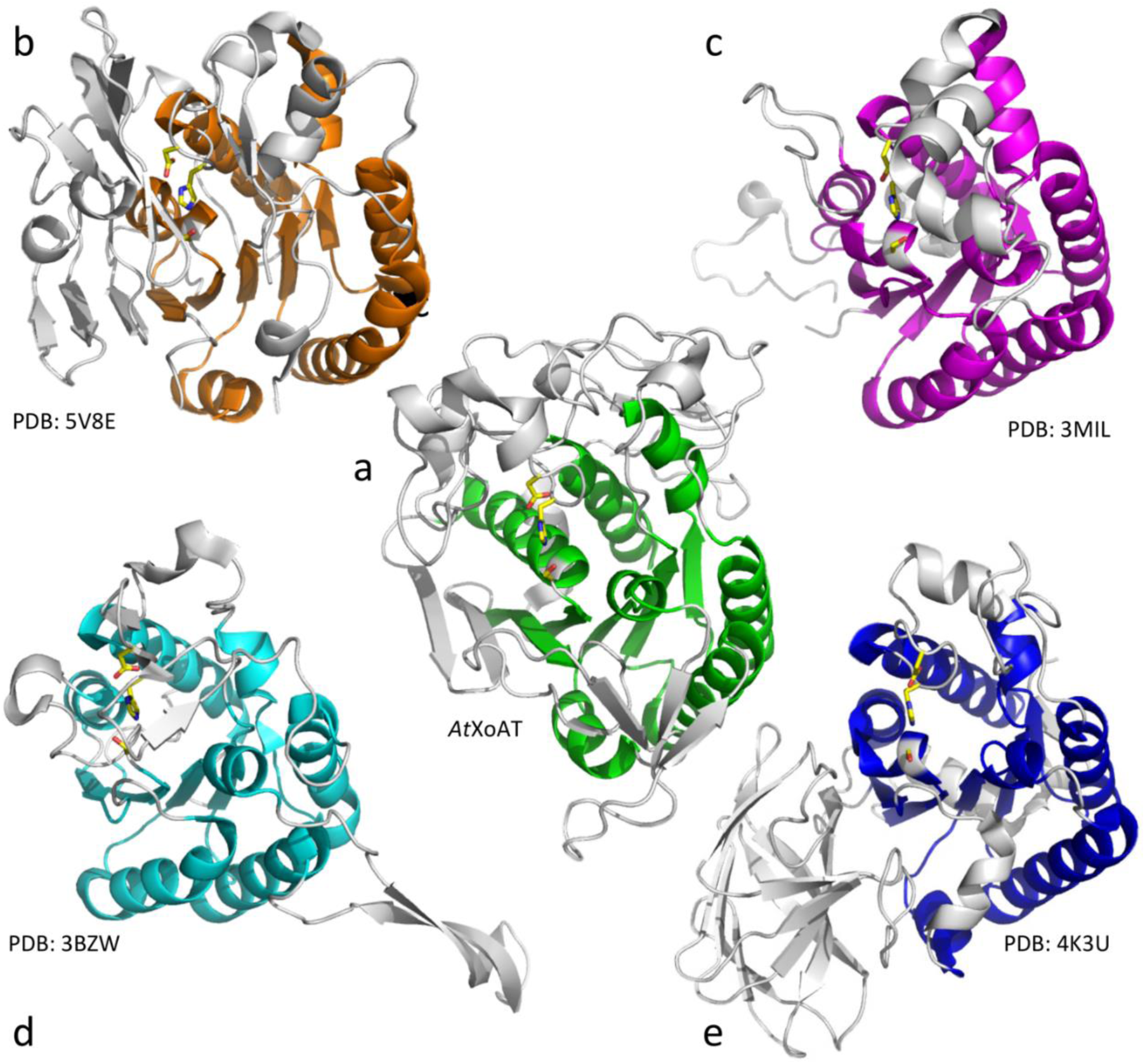
Active site and partial conservation of structural domains in A*t*XOAT1. Structural alignment of *At*XOAT1 (**A** - Center) with (**B** – Top Left) peptidoglycan *O*-acetyltransferase from *B. cerus* (PATB1) (PDB ID: 5V8E), (**C** – Top Right) Isoamyl acetate hydrolyzing esterase (PDB ID:3MIL), (**D** – Bottom Left) a putative lipase from *B. thetaiotamicron* (PDB ID 3BZW) and (**E** – Bottom right) peptidoglycan *O*-acetylesterase from *N. meningitidis* (PDB ID 4K3U). The secondary structure domains that demonstrate good alignment with *At*XOAT1 are depicted in solid colors while the non-aligned parts are greyed out. All structures share the Ser-His-Asp catalytic triad, which is shown in licorice representation with the carbons, oxygens and nitrogen atoms are colored yellow, red and blue, respectively.

### Mutagenic analyses implicate the roles key amino acids in *At*XOAT function

The C-terminal portion of *At*XOAT1 (AAs 193-481), comprised of the latter half of the plant-specific TBL (AAs 140-227) domain and the entire DUF231 domain (AAs 296-479), is categorized in the PFAM database as a domain from the GDSL/SGNH-like acyl-esterase family found in Pmr5 and Cas1p (PF13839), and shares two conserved motif blocks characteristic of the canonical GDSL/SGNH hydrolase superfamily (Fig. S7). As the name implies, enzymes of the SGNH-hydrolase family are typically classified based on the presence of four invariant residues (Ser-Gly-Asn-His) in conserved blocks (I, II, III, and V) that play important roles in specific aspects of catalysis. In these enzymes, the Ser in the GDS motif of block I and the Asp and His residues in the DxxH motif of block V form the catalytic triad, and the backbone amide of the Gly and side-chain amide of Asn residues in blocks II and III serve as hydrogen bond donors to stabilize the putative tetrahedral oxyanion intermediate (Akoh et al. 2004). Sequence analysis of the plant TBL family resulted in the identification of four highly conserved consensus regions; however, no conserved motifs were found representing SGNH blocks II or III (Fig. S7) or when compared across all plant species accessed from the Phytozome database (data not shown). The TBL and DUF231 domains contain the conserved GDSL and DxxH motifs of blocks I and V, respectively, that together form the putative Ser216-His465-Asp462 catalytic triad positioned at the bottom of the active site cleft (Fig. S7, Fig. 2). We observed that the position filled by Leucine in the GDS<u>L</u> motif can be occupied by any non-ring forming basic amino acid residue in plant TBL proteins. Directly downstream of the block I motif, there is a conserved RNQxxS motif, that we termed plant-specific TBL-block II. In plant TBL proteins, the block V **D**xx**H** motif is part of a larger **D**Cx**H**WCLPGxxD consensus region, which is highly conserved across all members of the family. We also identified a highly conserved, plant specific RxDxH motif that we termed plant-specific TBL-block III (Fig. S7). The catalytic region of *At*XOAT1 contains eight Cys residues that are 100% conserved, and half of these residues are present in the plant-specific, N-terminal PMR domain (PF14416), and all were found to all be involved in formation of the four disulfide bonds present in the smaller, less structured lobe of the enzyme.

To evaluate the involvement of these conserved amino acids in the TBL family that may play an important role in catalysis, we carried out site-directed mutagenesis of these residues in *At*XOAT1 by substituting them with alanine. As a control, we also mutated the catalytic residues (S216A, H465A and D462A), and verified that they abolish *O*-acetyltransferase activity as revealed by MALDI-TOF MS analysis of the reaction products (Fig. 4a). This confirmed that Ser216A, His465A and Asp462A are loss-of-function mutations that lack both transferase and esterase activity (Fig. 4b), and confirm the robustness of our *in vitro* assay. It is important to note that the esterase activity discussed here is indicated by release of *p*-nitrophenyl in the absence of acceptor, which occurs upon formation of the acyl-enzyme intermediate. Therefore, the combined assays allow us to measure both portions of the reaction: formation of the acyl-intermediate and transfer of the *O*-acetyl moiety from Ac-Ser216 to the xylosyl acceptor. Interestingly, when we incubated *At*XOAT1 with four electrophilic mechanism-based inhibitors of serine proteases, *N*-*p*-tosyl-L-phenylalanyl chloromethyl ketone (TPCK), 4-(2-aminoethyl)benzenesulfonyl fluoride hydrochloride (AEBSF), phenylmethylsulfonyl fluoride (PMSF) and methanesulfonyl fluoride (MSF), except for MSF, none of these protease inhibitors abrogated the catalytic activity of *At*XOAT1 as an *O*-acetyltransferase as acetylated products had been detected by MALDI-TOF MS after the reaction. Importantly, these data suggest that either *At*XOAT1 utilizes a different active site conformation compared to other serine proteases or that the majority of these compounds are unable to access the active site pocket (Fig. S8). A similar phenomenon was observed previously, in this case, PMSF was not able to completely inactivate *N. gonorrhoeae* PatB (Moynihan and Clarke 2014), while MSF abrogated both esterase and transferase activities of PatB in *Bacillus cereus*(Sychantha et al. 2018).

**Figure 4.**
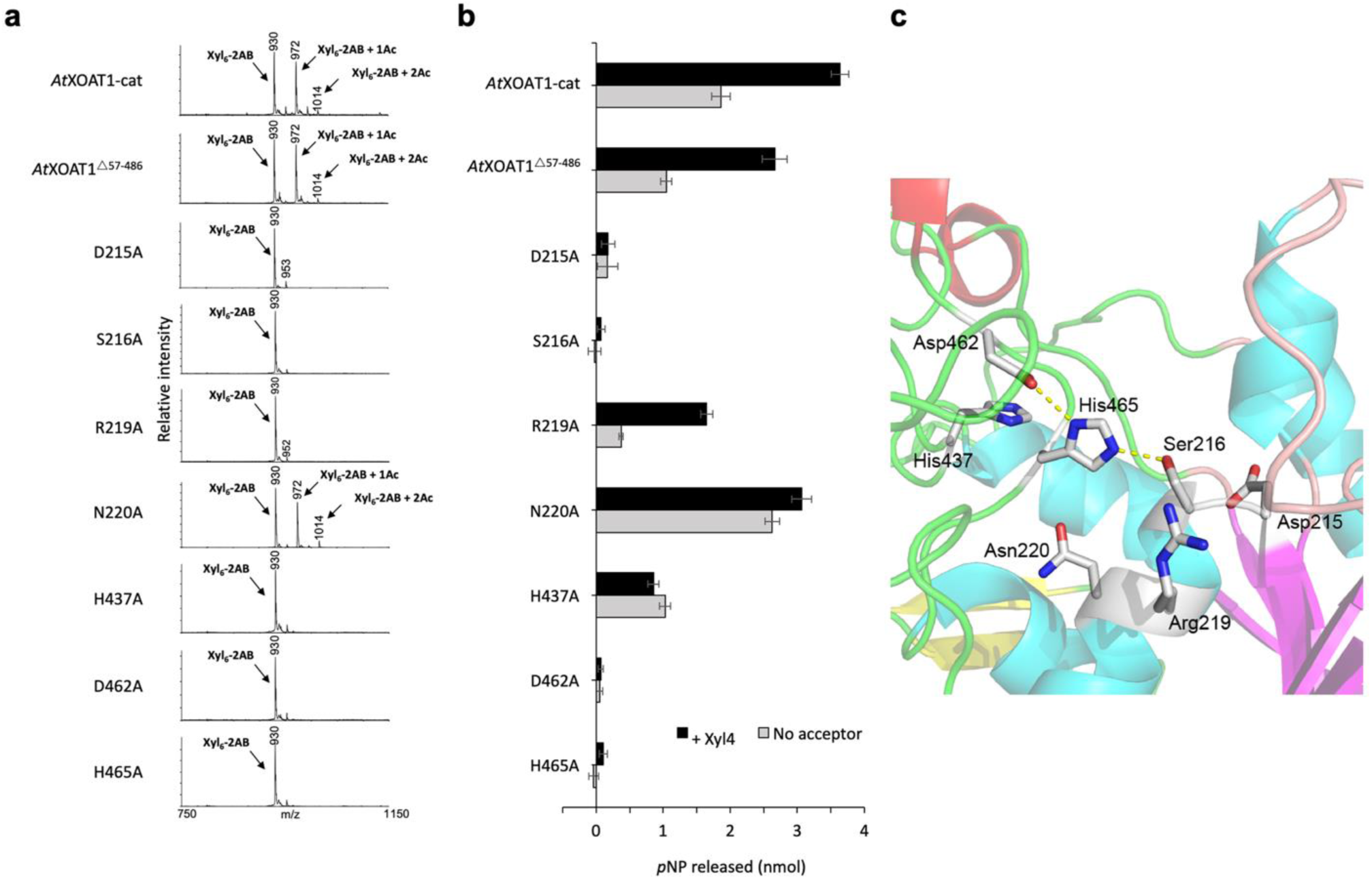
Enzymatic activity of *At*XOAT1 mutant variants compared with the wild type *At*XOAT1. (**A**) MALDI-TOF MS of the reaction products produced by *At*XOAT1 and its mutant variants. Each transfer of an acetyl group (Ac) increases the mass of the Xyl_6_-2AB acceptor by 42 Da as annotated [M+H]^+^ ions. (**B**) Comparative analysis of the esterase and transferase activities of of *At*XOAT1 and its variants determined by measuring the release of *p*NP from *p*NP-Ac (5 mM) formed in the absence (grey) or in the presence (black) of the acceptor substrate Xyl_4_ (5 mM), respectively. The error bars indicate mean ± s.d. from four independent assays. (**C**) Close-up view of the mutated residues at the *At*XOAT1 active site.

**Figure 5:**
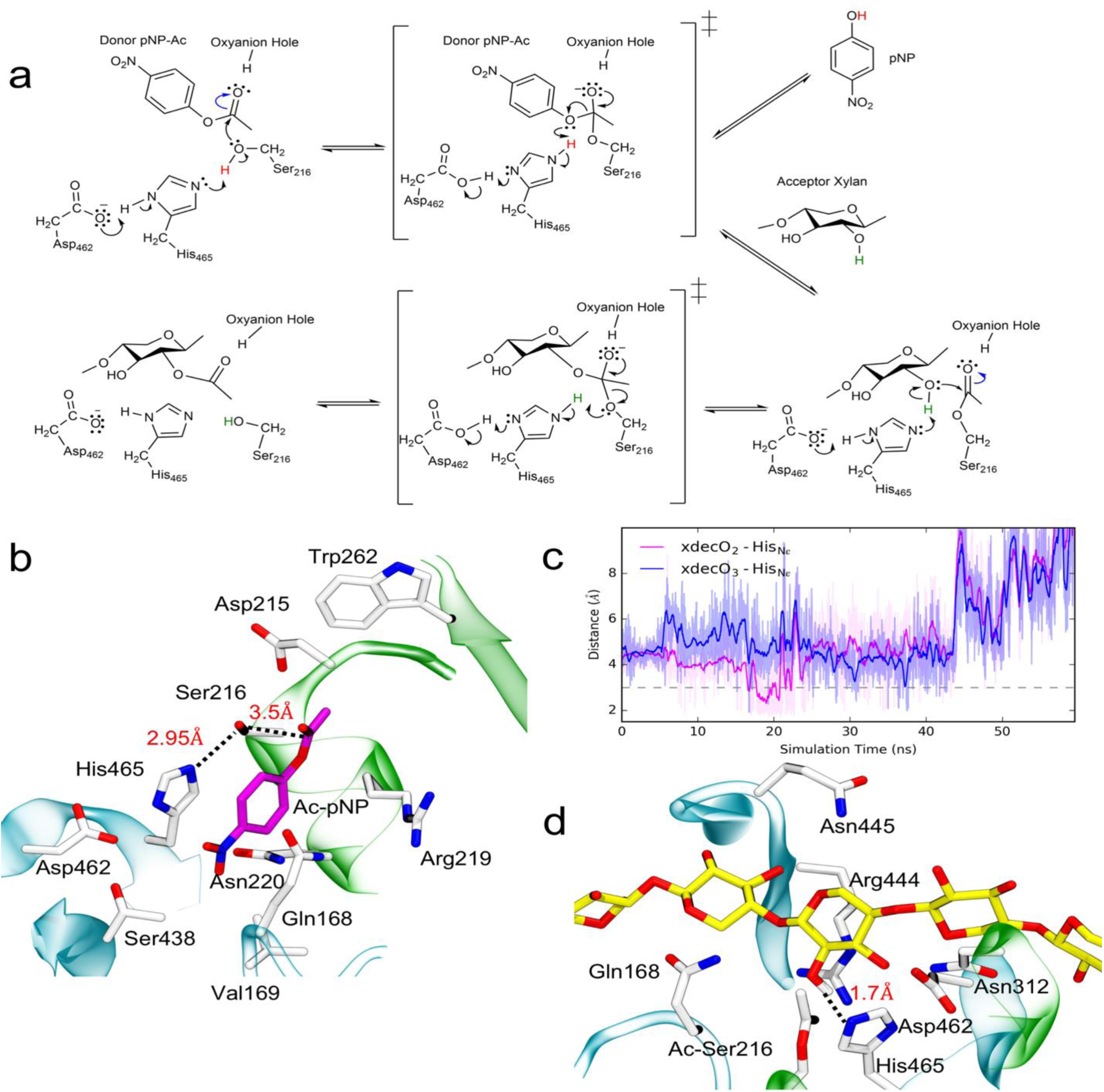
Reaction mechanism of *At*XOAT1. (**A**) Proposed reaction mechanism for acetylation of xylan oligosaccharides by *At*XOAT1. Rearrangement of the pi electrons within the His residue are not shown. (**B**) A snapshot from unbiased simulations of *At*XOAT1 in its donor bound state reveals catalytically competent configurations for the 1^st^ stage of the reaction mechanism. The donor molecule is shown in licorice representation with carbons colored magenta along with the important residues stabilizing it. Distances for the proton transfer and nucleophilic attack are shown with dashed lines. (**C**) Distances between protons on *O*-2 (magenta) and *O*-3 (blue) of the xylose ring closest to the catalytic triad indicate that the *O*-2 proton is consistently closer to the His465 Nitrogen. (**D**) A snapshot from an unbiased simulation of *At*XOAT1 in its acceptor (xylodecaose) bound state. Three xylose units closest to the active site are shown in licorice representation with the carbons colored yellow along with the important residues stabilizing it. The distance for proton transfer to generate the nucleophile on the xylose unit is shown with dashed lines.

An important factor for the transition state of the mechanism mediated by the Ser-His-Asp triad is the oxyanion hole that stabilizes the excess negative charge on the oxygen of the acetyl group. The backbone amide of the Gly and side-chain amide of an Asn residue in SGNH blocks II and III stabilize the putative tetrahedral oxyanion through hydrogen bonding in SGNH hydrolases (Akoh et al. 2004). However, residues that would be characteristic of blocks II and III that are important for formation of the oxyanion hole in the SGNH-hydrolase family are completely absent in the plant TBL family. A recent study suggested that Arg359 of the *O*-acetyltransferase PatB1 from *Bacillus cereus*, is a key component of the oxyanion hole in that enzyme (Sychantha et al. 2018). The conserved RNQxxS motif of TBL-block II contains an arginine (Arg219) that is correctly positioned in the active site and may function to stabilize the negative charges on the acetyl oxyanion during the formation of the tetrahedral reaction intermediates (Fig. 3c).

We designed an R219A variant of *At*XOAT1 to investigate its role in catalysis. The R219A variant is not capable of catalyzing the transfer of acetyl groups to Xyl_6_-2AB (Fig. 3a), indicating that Arg219 plays an important role in *At*XOAT1 catalysis. The hydrophilic side chain of Arg219 is positioned at the surface of the active site cleft, suggesting that this residue may play a role in forming the correct conformation of the catalytic site, and might be important for the oxyanion hole formation. Furthermore, Arg219 was observed to be an important active site residue involved in binding the donor molecule in the MD simulations. However, it is worth noting that even though transferase activity is abolished in the R219A variant as evidenced by MALDI-TOF MS of the reaction products (Fig. 3a), it retains esterase activity indicating that formation of the acyl-enzyme intermediate can still proceed to some extent (Fig. 3b). Furthermore, this esterase activity increases when xylo-oligo acceptors are present in the reaction, as revealed by the spectrophotometric assay (Fig. 3b), despite its inability to transfer an *O*-acetyl moiety to the acceptor (Fig. 3a). The increased esterase activity of R219A may indicate that the binding of the acceptor substrate stabilizes folding of *At*XOAT1 or contributes to formation of the oxyanion hole during the transfer of the acetyl group to the acceptor. Alternatively, in the absence of the guanidinium group of Arg219, the oligosaccharide may bind in an unproductive geometry that directs the movement of a water molecule to a position from which it can attack the serine-bound acetyl group, facilitating its hydrolysis. Thus, the binding of xylo-oligo substrates increases the esterase activity of R219A, and this mutation completely disrupts transferase activity, meaning Arg219 is critical for transferase activity and likely plays a role in limiting ester hydrolysis.

In most SGNH hydrolases, the side-chain amide of a conserved asparagine in SGNH block III is also important to maintain the stability of the oxyanion hole by serving as a hydrogen bond donor for stabilization of the transition state. While we hypothesize that the guanidinium moiety of an arginine residue in *At*XOAT1, like PatB1, is a key contributor to formation of the oxyanion hole, we also observe the presence of an asparagine residue (Asn220) that is located close to the Ser-His-Asp catalytic triad (Fig. 3c) – with its side chain amide pointing away from the active site. Thus, we evaluated the plausible role of Asn220; however, the N220A variant, despite spatial proximity, retained both acetyltransferase and esterase activity. At the end of the reaction, acetylated oligosaccharide products were detected by MALDI-TOF MS (Fig. 3a) and (deacetylated) *p*NP was detected spectrophotometrically (Fig. 3b), both in the presence and absence of acceptor substrate. This result indicates that the side chain amide of Asn220 is not involved in stabilizing the intermediate tetrahedral structure of the substrate during xylan *O*-acetylation. Interestingly, the N220A mutant has significantly more acetylhydrolase activity than either of the catalytically active controls, as indicated by the amount of *p*NP released in the absence of acceptor substrate (Fig. 3b). These data suggest that Asn220 might play a role in protecting the acyl-enzyme intermediate from hydrolysis, which would decrease the efficiency of the enzyme at low acceptor substrate concentrations.

Asp215 resides directly next to the catalytic Ser216, which is the site of nucleophilic attack during both stages of the mechanism (Fig. 3c). Mutation of Asp215 to Ala (D215A) abolished both *At*XOAT1’s esterase and transferase activities (Fig. 3a) as well as the level of released *p*NP in the hydrolysis assay was similar to that of the catalytically inactive control *At*XOAT1-nv (Fig. 3b). Since Asp215 lies very close to the catalytic site, it may contribute to stabilizing the active site conformation. Mutation of Asp215 resulted in a ∼75% decrease in production of soluble secreted protein (Table S1), indicating that this mutation may also lead to unstable protein folding, as observed previously with *At*FUT1 (Urbanowicz et al. 2017). The RxDxH motif we termed TBL-block III contains a highly conserved His437 residue that may play a role in substrate interactions. Interestingly, mutation of His437 to Ala (H437A) resulted in the production of no acetylated products based on detection using MALDI-TOF MS; however only 42% and 72% of hydrolysis activities retained in the H437A variant when reactions were respectively carried out with and without the acceptor substrate (Fig. 3b).

### Computational modelling enables insight into active site characteristics and reaction mechanism

Experimental efforts to obtain substrate bound structures of *At*XOAT1 with either a donor molecule (*p*NP-Ac or acetyl-CoA) or an acceptor (xylo-bi/tri/tetra/penta/hexa-oses) molecule were unsuccessful due to complications during the crystallization process (Supplementary Information Section I). Despite numerous attempts to crystallize the longer *At*XOAT1-fl construct, the only crystals that grew were formed by the *At*XOAT1-cat domain with no space left in the crystal lattice for the N-terminal variable region, even if it would have been disordered. We hypothesize that this tight packing interfered with our attempts to obtain a XOAT1-xylooligosaccharide co-complex. Given this lack of structural information on *At*XOAT1’s substrate bound states, gaining insight into the molecular details of the mechanism and mode of action required alternative strategies. Molecular simulations have previously been used to gain insight into substrate binding poses and the catalytic mechanism in cases where experimental characterization of enzyme-substrate complexes has been unsuccessful (Bharadwaj et al. 2013)(22). As outlined in the following paragraphs, molecular simulations reveal *At*XOAT1’s putative active site and the ability of its substrate binding groove to stabilize acceptor and donor substrates.

It has also been experimentally shown that *At*XOAT1 demonstrates increased activity with longer xylan oligosaccharides (>DP3) and catalyzes the addition of multiple acetyl moieties onto a single acceptor necessitating an active site that can accommodate larger acceptor oligosaccharides (Fig. S2). MD simulations of *At*XOAT1 in the *apo* state reveal significant flexibility in loops surrounding the putative substrate binding groove. The presence of a Ser-His-Asp triad at the base of this loop further strengthens the hypothesis that the cleft is an extension of the active site to enable binding oligosaccharide substrates. By analogy to serine proteases and esterases (Akoh et al. 2004), the Ser residue acts as a nucleophile assisted by the His residue, which acts as a base whose excess positive charge is stabilized by a hydrogen bond with the Asp residue. Figure S9 depicts the two key hydrogen bonds that are necessary for the proper functioning of this catalytic triad. In the *apo* state, it is observed that the acid-base distance (Asp462 O to His465 H) is within 2 Å throughout the MD simulation, whereas the base-nucleophile distance is frequently (22% of the simulation time) within 3 Å. It must be noted that the Asp462 and His465 are a part of the same minor lobe of *At*XOAT1, whereas the nucleophile is situated on an α-helix that is part of the major lobe of *At*XOAT1 (Fig. 2c). This may explain the observation that the acid-base distances are shorter than the base-nucleophile distances.

Separate MD simulations of *At*XOAT1 in the donor and acceptor bound states revealed the ability of the active site and the putative binding groove to accommodate the surrogate donor molecule (*p*NP-Ac) and the acceptor xylodecaose substrate over simulation times in excess of 50ns. These observations of substrate bound configurations from unbiased MD simulations combined with data from mutagenesis experiments enables insight into *At*XOAT1’s catalytic mode of action. Structural alignments (Section 2.2) of sub-regions of the *At*XOAT1 crystal structure with other proteins revealed conservation of important structural domains; as well as the putative catalytic triad amongst these enzymes. Furthermore, MD simulations of *At*XOAT1 in the *apo*, donor- and acceptor-bound states established the stability of the catalytic triad; as well as the ability of the active site to bind the acceptor and donor molecules. The mutagenesis experiments further substantiated the indispensable role of the Ser216-His465-Asp462 catalytic triad and helped identify residues that may contribute to formation of an oxyanion hole in *At*XOAT1. Based on these observations we propose that XOAT1 utilizes a double displacement bi-bi reaction to catalyze xylan 2-*O*-acetylation consisting of two stages as illustrated in Figure 4a. The first stage of catalysis involves acetylation of Ser216 upon binding of the acetyl donor, in this case the surrogate donor *p*NP-Ac is depicted. Coordination of His465 with the unprotonated Asp462 allows His465 to act as a general base, facilitating the nucleophilic attack of the acetyl group of the donor by Ser216. Arg219, contributes to formation of an oxyanion hole that stabilizes the putative tetrahedral transition state. The transfer (via His465) of the Ser216-proton (red font) onto the phenolic oxygen of the donor facilitates release of the deacylated donor (*p*-nitrophenol as shown) and acetylated Ser216 (acyl-enzyme intermediate), which is supported by the LC-MS/MS data (Fig. 1d, Fig. S3 and Fig. S4) showing that *At*XOAT is able to utilize all acetyl donors investigated in formation of the acyl-enzyme intermediate.

## Dicussion

Unraveling the identity, selectivity, and catalytic mechanisms of plant biocatalysts involved in cell wall formation is vital to understanding the roles these enzymes play in generating structures critical to the composite nature of the plant cell wall. Xylan, the second most abundant component of plant biomass after cellulose, forms an integral part of the wall and defines its architecture and structural properties. An increased understanding of gene pathways involved in xylan synthesis provides insight into the mechanism of cell wall formation, which is fundamental to exploiting wall structural plasticity in agronomically productive crops. Enzymes that catalyze the addition of substituents, such as acetyl groups to the xylan backbone (e.g., *At*XOAT1), are of significant interest as they play key roles in shaping the polysaccharide’s physical properties. The importance of this single enzyme in plant growth and development is substantiated by the fact that mutation of *At*XOAT1 in Arabidopsis results in plants with a 40% reduction in xylan *O*-acetylation that exhibit collapsed xylem vessels, and manifest several pleiotropic phenotypes related to stress responses (Pauly and Ramírez 2018).

Expression of the soluble catalytic domain of *At*XOAT1 (*At*XOAT-cat, *At*XOAT^133-487^) for structural analysis was achieved by transient transfection of suspension culture HEK293S cells lacking *N*-acetylglucosaminyltransferase I (GnTI-) using a fusion protein strategy similar to our prior studies on mammalian GTs (Moremen et al. 2018) and *Arabidopsis thaliana* fucosyltransferase 1 (*At*FUT1) (Urbanowicz et al. 2017). In this study, we demonstrate the successful expression, purification and crystallization of AtXOAT1 from *Arabidopsis thaliana*. The catalytic domain of *At*XOAT1 (AtXOAT-cat, AtXOAT^133-487^) adopts a unique structure, bearing some similarities the α/β/α topology of members of the GDSL-like lipase/acylhydrolase family, and is comprised of three β-sheets composed of seven, four and two β-strands, nine α-helices and three α-helical turns. A deep cleft divides the molecule into two unequal lobes that open to accommodate the mono *O*-acetylation of linear β-1,4-xylan and contains a Ser216-His465-Asp462 catalytic triad at its base. Real time NMR experiments confirm AtXOAT1 specifically transfers *O*-acetyl moieties specifically to the 2-position of the xylan backbone. Further, this technique was applied to study acetyl migration in real time, and showed that the *O*-acetyl at the 2-position migrates spontaneously to the 3-position in the absence of enzyme. A double displacement bi-bi mechanism involving the Ser216-His465-Asp462 triad is proposed and substantiated by both molecular simulation results and biochemical experiments confirming the formation of the acyl-enzyme intermediate, Ac-Ser216. In addition, we propose that *At*XOAT1 likely unconventionally employs an arginine (Arg219) residue to conduct the role of an oxyanion hole in the mechanism.

These mechanistic insights into an enzyme involved in plant polysaccharide acetylation and the biosynthesis of the predominant hemicellulosic polysaccharide xylan represent a major leap forward in understanding plant cell wall biochemistry. This work lays the basis for further studies exploring the nature of xylan acetylation, including the interplay between glycosyl and non-glycosyl substituent patterning on the glycopolymer backbone and the regiospecificity of acetylation and the molecular level impacts that this modification has on xylan interactions with other cell wall components and ultimately plant cell wall properties.

## Materials and Methods

### Experimental Design

The primary objective of this study was to use a multidisciplinary approach to understand the enzymatic mechanism underlying the acetylation process of xylan in plants. Specifically, the promiscuity of *At*XOAT1 to accommodate different acetyl donor substrates was first analyzed through the *in vitro* assays followed by MALDI-TOF MS, and the regiospecificity of *At*XOAT1 was then unambiguously distinguished by real-time 1H NMR spectroscopy. The formation of an acyl-enzyme intermediate involving a Ser-His-Asp catalytic triad was further confirmed by LC-MS/MS analysis. Moreover, we elucidated *At*XOAT1’s structure using X-ray crystallography, following by investigating the active site dynamics and the substrate binding modes using molecular dynamics (MD) simulation. The viable mechanism revealed by computational modeling was further validated through mutagenesis experiment including *in vitro* acetyltransferase assays and chromatic hydrolysis assays of the *At*XOAT1 variants.

### Expression and purification of fusion proteins

Fusion protein expression and purification was accomplished as described previously (Urbanowicz et al. 2014). Briefly, total RNAs extracted from inflorescence stems of wild-type *A. thaliana* (Col-0) were used to generate cDNAs using RevertAid First Strand cDNA Synthesis Kit (Thermo Scientific), and the sequences of the truncated coding-region of XOAT1 were then amplified from the cDNAs by using the following primer pairs: TBL29/ESK1, 5’-AACTTGTACTTTCAAGGCTCAAAACCTCACGACGTC-3’ and 5’-ACAAGAAAGCTGGGTCCTAACGGGAAATGATACGTGT-3’. The resulting PCR product was cloned into a mammalian expression vector, pGEn2-DEST(Moremen et al. 2018), as described previously (22). The expression plasmids were purified and transformed into HEK cells (FreeStyle™ 293-F cell line, Life Technologies) as previously described(Moremen et al. 2018).

Recombinant proteins were secreted into the culture media by HEK293 cells, and were purified using HisTrap HP column and ÄKTA protein purification system (GE Healthcare Life Sciences). For crystallization ESK1 samples were further purified using Source 15Q anion exchange chromatography with 20 mM Tris pH 7.5 and 0 to 0.35 M NaCl gradient. Finally, Superdex 75 pg (26/60) size exclusion chromatography column in 20 mM Tris pH 7.5, 100 mM NaCl was used to complete the purification. SDS-PAGE followed by Coomassie Brilliant Blue R-250 (Bio-Rad) staining was carried out to confirm the purity of the proteins. For the activity assays, proteins were dialyzed in HEPES sodium salt-HCl (75 mM, pH 7.0) with Chelex-100 resin (5 g/L, Bio-Rad) to remove any divalent metal contaminants. After dialysis, the proteins were concentrated using Amicon Ultra centrifugal filter devices (30 kDa molecular weight cut-off, EMD Millipore), and were quantified by Pierce™ BCA Protein Assay Kit (Thermo Scientific). The resulting fusion proteins comprise NH_2_-terminal signal peptides, 8×His-tags, AviTag recognition sites, Green fluorescent proteins, TEV protease recognition sites, and the catalytic domains of the corresponding enzymes.

Mutated variants of *At*XOAT1 were generated by site-directed mutagenesis using the Q5® Site-Directed Mutagenesis Kit (New England Biolabs) according to the manufacturer’s instructions using pGEN2-DEST-*At*XOAT1 as a template. Oligonucleotide primers used to generate the base changes and expression levels of the wild-type and mutant variants are listed in Table X. The introduction of mutations was confirmed by sequencing (Eurofins, USA).

### Crystallization

*At*XOAT1-cat crystals were initially obtained with sitting drop vapor diffusion using a 96-well plate with PEG ion HT screen from Hampton Research (Aliso Viejo, CA). Fifty µL of well solution was added to the reservoir and drops were made with 0.2 µL of well solution and 0.2 µL of protein solution using a Phoenix crystallization robot (Art Robbins Instruments, Sunnyvale, CA). The crystals were grown at 20°C using an optimization screen containing 0.1 M MES pH 6.0 to 7.0 and 15% to 22% w/v PEG 3350, 0.02 M Calcium chloride dihydrate, 0.02 M Cadmium chloride hydrate and 0.02 M Cobalt(II) chloride hexahydrate. The protein solutions contained 5.62 mg/mL of protein, 20 mM Tris pH 7.5, 100 mM NaCl and 20 mM 4-Nitrophenyl acetate. Crystals were soaked in well solution with additional 10% glycerol and 5% ethylene glycol for additional cryo protection.

### Data collection and processing

The *At*XOAT1-cat crystals were flash frozen in a nitrogen gas stream at 100 K before home source data collection using an in-house Bruker X8 MicroStar X-Ray generator with Helios mirrors and Bruker Platinum 135 CCD detector. Data were indexed and processed with the Bruker Suite of programs version 2014.9 (Bruker AXS, Madison, WI).

### Structure solution and refinement

Intensities were converted into structure factors and 5% of the reflections were flagged for R_free_ calculations using programs F2MTZ, Truncate, CAD and Unique from the CCP4 package of programs (Winn et al. 2011). Crank2 (Skubak and Pannu 2013) was used to solve the structure utilizing sulfur Single-wavelength anomalous dispersion (Hendrickson and Teeter 1981). Refinement and manual correction was performed using REFMAC5 (Murshudov et al. 2011) version 5.8.135 and Coot (Emsley et al. 2010) version 0.8.2. The MOLPROBITY method (Chen et al. 2010) was used to analyze the Ramachandran plot and root mean square deviations (rmsd) of bond lengths and angles were calculated from ideal values of Engh and Huber stereo chemical parameters (Engh and Huber 1991). Wilson B-factor was calculated using CTRUNCATE version 1.15.10 (Winn et al. 2011). The data collection and refinement statistics are shown in Table I.

**Table I:**
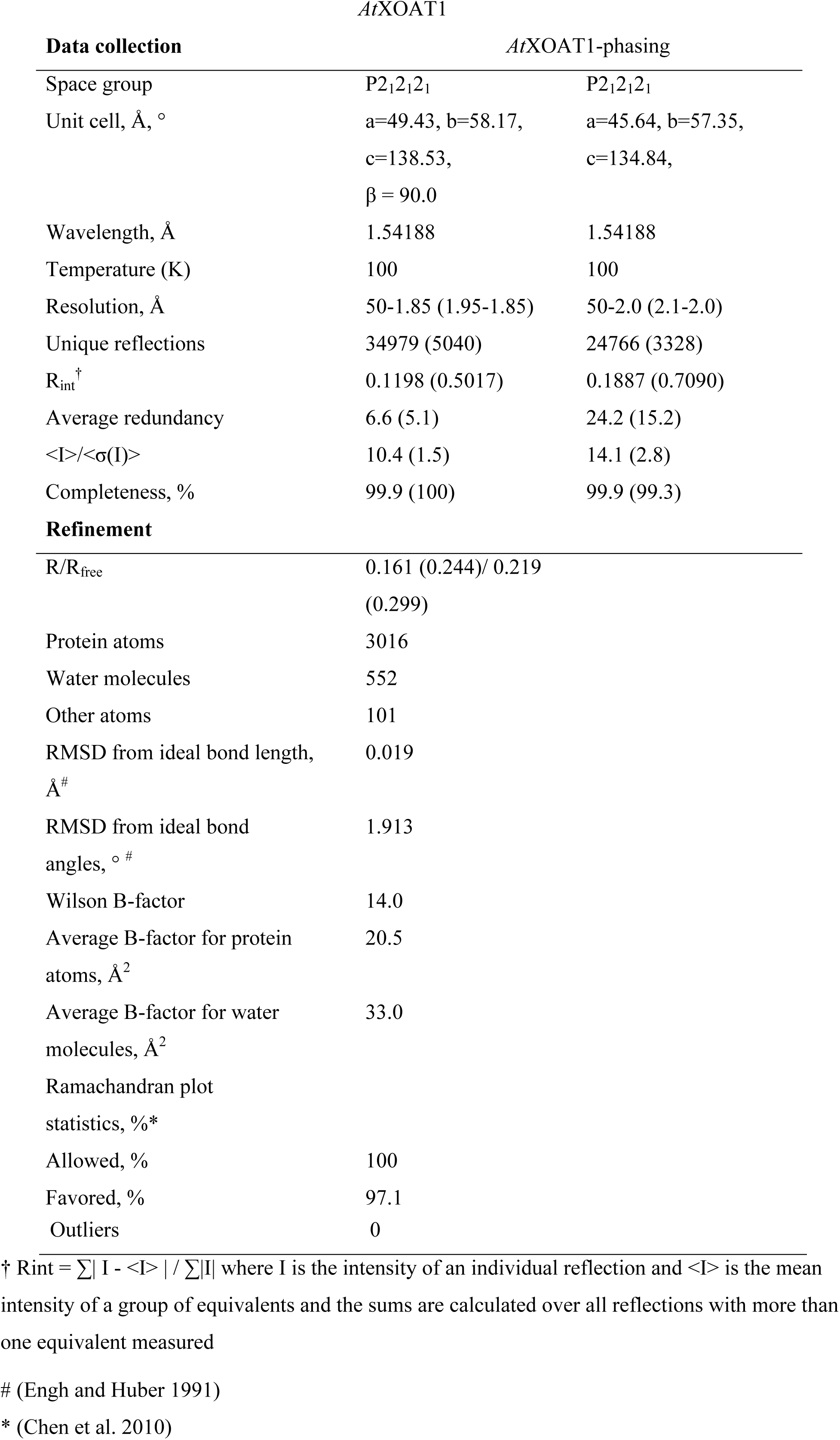
X-ray data collection and refinement statistics. Statistics for the highest resolution bin are shown in parenthesis.

### Donor and acceptor substrate specificities of AtXOAT1

To determine the activity of acetyltransferase on different donor substrates, the standard assays (15 µL) were performed using acetylsalicylic acid (1 mM), acetyl-CoA (1 mM), or *p*NP-Ac (1 mM) as donors, and Xyl_6_-2AB (0.25 mM) as acceptor substrates in HEPES sodium salt-HCl (75 mM, pH 6.8) with purified proteins (4 µM). The reactions were incubated at room temperature for indicated periods of time, and the products were then analyzed by Matrix-assisted laser desorption/ionization mass spectrometry (MALDI-TOF MS) using a Microflex LT spectrometer (Bruker) as described previously (22). Briefly, aliquots (5 µL) of the reactions were mixed with Dowex-50 cation exchanger resin (1 µL suspension in water), and the mixture was incubated at room temperature for 30 min. Resin was pelleted by centrifugation and 1 µL of the supernatant was directly mixed with 1 µL of DHB matrix solution (20 mg/mL 2,5-dihydroxybenzoic acid in 50% MeOH), on a stainless steel MALDI target plate then concentrated to dryness using a hair dryer. Each positive ion spectrum was generated by summation of 200 laser shots.

Xylooligosaccharides (Xyl_2_, Xyl_3_, Xyl_4_, Xyl_5_ and Xyl_6_) were purchased from Megazyme. The acceptor substrate specificities were determined by the similar assays mentioned above but with increased amount of acetylsalicylic acid (5 mM) as the donor and non-labeled xylo-oligosaccharides (3 mM), Xyl_2_, Xyl_3_, Xyl_4_, Xyl_5_ and Xyl_6_, as the acceptor substrates, and acetylated products were analyzed by MALDI-TOF MS. On the other hand, the acetyltransferase activity of *At*XOAT1 on xylo-oligosaccharide was examined by the hydrolysis assays using *p*NP-Ac (5 mM) as the donor, and xylopentaose (Xyl_5_) with different concentrations (0, 1, 5 and 25 mM) as the acceptor substrate in HEPES sodium salt-HCl (75 mM, pH6.8) with purified proteins (5 µM). The reactions were incubated for an hour, and the hydrolysis of *p*NP-Ac in the reaction was determined by measuring the absorbance of the released *para*-nitrophenol (*p*NP) at 350 nm, using Epoch Microplate Spectrometer (BioTek®) every minute during the reaction period.

### Determination of catalytic activity of AtXOAT1 constructs and variants

To evaluate the *O*-acetyltransferase activities of different *At*XOAT1 constructs and its mutated variants, the activity assays were performed by incubating the purified enzymes (4 µM) with acetylsalicylic acid (1 mM) and Xyl_6_-2AB (0.25 mM) in HEPES sodium salt-HCl (75 mM, pH 6.8) for 2 h. The acetylated products were analyzed by MALDI-TOF MS as described previously. To further investigate whether *At*XOAT1 variants possess xylan *O*-acetyltransferase and/or *O*-acetylesterase activities, assays were carried out by incubating the enzymes (5 µM) with or without non-labeled Xyl_4_ (5 mM) as the acceptor and *p*NP-Ac (5 mM) as the donor substrate for 60 min. The amount of released *p*NP in the reaction was quantified by measuring the absorbance at 350 nm using Epoch Microplate Spectrometer (BioTek®) as the same described above. The non-catalytic truncated protein containing only the N-terminal variable region and lacking the catalytic domain, AtXOAT1-nv (Fig. S5b and c), was used as a control. Each transferase and esterase activity of the AtXOAT1 variants was obtained by subtracting the amount of *p*NP released in the control sample containing AtXOAT1-nv from each standard reaction containing different AtXOAT1 variants with or without the acceptor.

### Peptide analysis of acyl-AtXOAT1 by LC-MS/MS

I*n vitro* assays were carried out by incubating *At*XOAT1 (4 µM) with four different acetyl donors (1 mM): acetyl-CoA, acetyl salicylic acid, *para*-nitrophenyl acetate and 4-methylumbelliferyl acetate in HEPES sodium salt buffer (pH 6.8; 75 mM) for 4 min at room temperature. The reactions were then stopped immediately by flash freezing in a dry ice/ethanol cooling bath (−78 °C). All the donor substrate stocks were prepared in DMSO, and the final percentage of DMSO in all reactions was 5% (v/v). Control reactions were identical, but no donor substrate was added.

The prepared samples were reduced, alkylated, and then digested with trypsin (sequencing grade; Promega) in Tris-HCl (50mM, pH 8.2) buffer overnight. After protease digestion, samples were desalted and filtered prior to LC-MS analysis. NanoLC-MS/MS was performed on an Orbitrap Fusion Tribrid Mass Spectrometer (ThermoFisher Scientific) coupled to an Ultimate3000 RSLCnano low flow liquid chromatography system (ThermoFisher Scientific), and equipped with a nanospray ion source. Prepared peptide samples were injected onto a loading column (ThermoFisher Scientific, C18, volume 2µL) before being transferred to the separation column (Acclaim PepMap 100 C18 – 0.075×150mm with 3µm particle size). The separation was performed under a linear gradient from 5% to 100% B (80% Acetonitrile, 0.1% formic acid) at a flow rate of 0.3µL/min. The analysis was run in automatic mode collecting an MS scan (full FTMS at 150-2000 m/z; resolution 120 000) followed by data dependent MS/MS scans (CID) at a cycle time of 3 sec. The data was manually processed using QualBrowser (ThermoFisher Scientific).

### RT-NMR analysis of acetyl group migration

To verify the positional specificity of AtXOAT1-cat, the reaction of *O*-acetylation of Xyl6 was monitored using real-time 1H NMR spectroscopy. 200 µg Xyl_6_ was mixed with 4 mM Ac-CoA and 9 µM purified enzyme in 100 mM potassium phosphate buffer (pH 6.8) in D_2_O (99.9%; Cambridge Isotope Laboratories), and the reaction (110 uL total volume) was rapidly transferred to the NMR spectrometer. Previously, we employed potassium bicarbonate as a buffer in our real-time NMR experiments (18); however, biocarbonate buffers are unstable and become alkaline upon exposure to atmospheric CO_2_, increasing the rate of acetyl migration in solution. Due to this observation, we have now optimized our real-time NMR conditions and used phosphate buffer, which is a more stable buffer. Data acquisition started 10 min after the reaction was initiated, and data were recorded at 298 K with an Agilent-NMR spectrometer operating at 600 MHz equipped with a 5-mm NMR cold probe. The 1D ^1^H spectra consist of 16 transients and were acquired with water presaturation every 30 min over a 20-hour period. The spectral array generated represents the reaction progress. The amount of acetylated xylosyl residues generated during the reaction were quantified by integrating the resonance peaks corresponding to the methyl protons of the acetyl groups attached to *O*-2 and *O*-3 of the xylosyl residues, and the calculation was based on the initial concentration of Ac-CoA added into the reaction. Data processing was performed using MestReNova software (Mestrelab Research S.L., Universidad de Santiago de Compostela, Spain).

Non-enzymatic acetyl group migration was monitored in real-time by ^1^H NMR. To generate an adequate amount of acetylated Xyl_6_ that meets the sensitivity of the NMR spectrometer, scaled up reactions were performed in parallel to those prepared for real-time NMR analysis of catalysis. Briefly, acetylated Xyl_6_ was generated by incubating AtXOAT1 (9 µM) with non-labeled Xyl6 (200 µg) and Ac-CoA (4 mM) in 100 mM potassium phosphate buffer in D2O (99.9%; Cambridge Isotope Laboratories) for 6 h at room temperature. The enzyme was then removed via diafiltration by passing the mixture through a 30 kDa centrifugal filter device (Amicon®, Merck Millipore Ltd., Ireland). The absence of AtXOAT1 in the filtrate, and its complete retention in the filter device, was confirmed by both SDS-PAGE using a precast Mini-PROTEIN® TGX Stain-Free™ Gel (Bio-Rad Laboratories, Inc.) and by measuring the direct absorbance at 280 nm using a NanoDrop™ Spectrophotometer (Thermo Fisher Scientific). The filtrate was immediately transferred to the NMR spectrometer, and the signals that correspond to the CH_3_ of the *O*-acetyl substituents at different positions were monitored by arrayed ^1^H NMR according to the following process. Data were recorded at 298 K with an Agilent-NMR spectrometer equipped with a 5-mm NMR cold probe operating at 600 MHz. Each 1D ^1^H spectra consists of 16 transients, and were acquired with water presaturation every 30 min for 14 hrs overnight. The spectral array generated represents the reaction progress. Samples were never subjected to lyophilization.

### *In vitro* assays of AtXOAT1 with protease Inhibitors

The *in vitro* assays with protease inhibitors were carried out as the standard assays mentioned previously but with addition of 80 μM *N*-*p*-tosyl-L-phenylalanyl chloromethyl ketone (TPCK), 0.8 mM 4-(2-aminoethyl)benzenesulfonyl fluoride hydrochloride (AEBSF), 4 mM phenylmethylsulfonyl fluoride (PMSF) or 5 mM methanesulfonyl fluoride (MSF) in the reaction. After overnight incubation, the acetylated products were analyzed by MALDI-TOF MS.

### Computational Modeling

Docking and molecular dynamics simulations have been successful in gaining insight into plausible substrate binding poses and identify the important active site residues that aid substrate binding.(Bharadwaj et al. 2013; Urbanowicz et al. 2017) AutoDock4 and AutoDockTools4 software package was used for the docking the acceptor (xylodecaose) and donor (*p*NP-Ac) molecules to the 1.85 Å *At*XOAT1 structure(Morris et al. 2009). Plausible binding poses were identified as starting points for molecular dynamics simulations. The four disulfide bonds identified in the crystal structure, between residues 465-386, 469-173, 229-164 and 142-193 were incorporated and the protonation states of titratable amino acid residues was estimated using the H++ web server.(Anandakrishnan et al. 2012) Fully solvated systems were generated for the protein in its apo and bound states. The CHARMM36 forcefield was used for the protein(Huang and MacKerell 2013), xylodecaose substrate(Guvench et al. 2009), the CGENFF force field for pNP-Ac(Vanommeslaeghe et al. 2010), and the TIP3P forcefield for modelling water(Jorgensen et al. 1983). Three systems of *At*XOAT in different states were built - (i) apo state (no substrate bound) (ii) donor bound state (*At*XOAT1 with six *p*NP-Ac molecules bound) and (iii) acceptor bound state (*At*XOAT1 with acetylated Ser216 and a xylodecaose substrate bound). The CGENFF protocol was used to obtain parameters for the acetylated serine residue in the acceptor bound state.(Vanommeslaeghe et al. 2010) The system set-ups for the bound states was obtained based on preliminary docking studies and hence the simulation procedure for the protein-substrate complexes involved multiple minimizations and short equilibration runs with restraints on the substrate. The short equilibration runs involved sequential runs starting with a fixed substrate, followed by restraints on the substrate which were gradually released for unrestrained production runs. The production runs for data analysis involved 60 ns unbiased runs with all H-atoms restrained using the SHAKE algorithm, a 2 fs time step, periodic boundary conditions, a non-bonded cutoff of 11 Å, the particle mesh Ewald for long range electrostatics and frames saved every 100 ps.(Darden et al. 1993; Ryckaert et al. 1977) All MD simulations utilized the domdec MD engine that is part of the CHARMM MD package (Brooks et al. 2009; Hynninen and Crowley 2014). The trajectories were analyzed for root mean square fluctuations, interaction energies, catalytically important distances and volumetric occupancies. The RMSF and distance calculations were performed with pytraj (python version of CPPTRAJ (Roe and Cheatham 2013)) and the volumetric occupancies were calculated using the volmap tool in VMD (Humphrey et al. 1996).

### Statistical Analysis

All data are present as mean ± SD except for Fig. S1 (the quantitative data of the duplicates and their average are displayed). Statistical significance was determined with the two-tailed Student’s *t*-test through Microsoft® Excel Version 15.32 (2017). *, *p* < 0.05; **, *p* < 0.01. The number of replicates is provided in the figure legend for each experiment.

## Accession Numbers

The atomic coordinates of XOAT1cat have been deposited into the Protein DataBank (accession code 6CCI; http://www.rcsb.org/pdb). Sequence data for the genes that were described in this article can be found in the Arabidopsis Genome Initiative or GenBank/EMBL databases under the following Arabidopsis AGI locus identifiers: At5g06700 (AtTBR); At3g12060 (AtTBL1); At1g60790 (AtTBL2); At5g01360 (AtTBL3/AtXOAT4); At5g49340 (AtTBL4); At5g20590 (AtTBL5); At3g62390 (AtTBL6); At1g48880 (AtTBL7); At3g11570 (AtTBL8); At5g06230 (AtTBL9); At3g06080 (AtTBL10); At5g19160 (AtTBL11); At5g64470 (AtTBL12); At2g14530 (AtTBL13); At5g64020 (AtTBL14); At2g37720 (AtTBL15); At5g20680 (AtTBL16); At5g51640 (AtTBL17/AtYLS7); At4g25360 (AtTBL18); At5g15900 (AtTBL19); At3g02440 (AtTBL20); At5g15890 (AtTBL21); At3g28150 (AtTBL22/AtAXY4L); At4g11090 (AtTBL23/AtMOAT1); (AtTBL24/AtMOAT2); At1g01430 (AtTBL25/AtMOAT3); At4g01080 (AtTBL26/AtMOAT4);At1g70230 (AtTBL27/AtAXY4);At2g40150 (AtTBL28/AtXOAT2); At3g55990(AtTBL29/AtXOAT1/AtES K1); At2g40160 (AtTBL30/AtXOAT3); At1g73140 AtTBL31/AtXOAT5); At3g11030 (AtTBL32/AtXOAT6); At2g40320 (AtTBL33/AtXOAT7); At2g38320 (AtTBL34/AtXOAT8); At5g01620 (AtTBL35/AtXOAT9); At3g54260 (AtTBL36); At2g34070 (AtTBL37); At1g29050 (AtTBL38); At2g42570 (AtTBL39); At2g31110 (AtTBL40); At3g14850 (AtTBL41); At1g78710 (AtTBL42); At2g30900 (AtTBL43); At5g58600 (AtTBL44/AtPMR5); At2g30010 (AtTBL45).

## Supplemental Data

Supplemental Materials and Methods. Supplementary description of crystallization trials and structure determination

Table S1. Primer sequences for gene cloning and site-directed mutagenesis for generation of the expression constructs

Figure S1. Activity curves of *At*XOAT1 as an *O*-acetyltransferase.

Figure S2. MALDI-TOF MS of the acetylated xylo-oligosacharides generated by *At*XOAT1.

Figure S3. Quantitation of the XOAT1 acyl-enzyme intermediate.

Figure S4. MS/MS Fragmentation Spectra (HCD) of m/z corresponding to the acetylated peptide MMFVGDSLNR (m/z 606.2840 z=2), where S indicates the site of modification.

Figure S5. Domain architectures of the designed expression constructs of *At*XOAT1 and their corresponding *O*-acetyltransferase activities.

Figure S6. Protein topology diagram for AtXOAT1-cat.

Figure S7. Sequence alignment of the *Arabidopsis* TBL protein family.

Figure S8. MALDI-TOF MS of the reaction products produced by *At*XOAT1 with serine protease inhibitors.

Figure S9. Stability of the Catalytic Triad.

## Acknowledgements

Funding was provided in part by the BioEnergy Science Center (BESC) and the Center for Bioenergy Innovation (CBI), from the U.S. Department of Energy Bioenergy Research Centers supported by the Office of Biological and Environmental Research in the DOE Office of Science. This work was also supported by the Laboratory Directed Research and Development (LDRD) Program at NREL. This work was authored in part by Alliance for Sustainable Energy, LLC, the Manager and Operator of the National Renewable Energy Laboratory for the U.S. Department of Energy (DOE) under Contract No. DE-AC36-08GO28308. The views expressed in the article do not necessarily represent the views of the DOE or the U.S. Government. The U.S. Government and the publisher, by accepting the article for publication, acknowledges that the U.S. Government retains a nonexclusive, paid-up, irrevocable, worldwide license to publish or reproduce the published form of this work, or allow others to do so, for U.S. Government purposes. This research was also supported by NIH grants P41GM103390, P01GM107012 and 1S10OD018530.

## Author Contributions

V.V.L., H.T.W., V.S.B., P.M.A., M.J.P., J.Y.Y., S.A.A-H., Y.J.B and B.R.U designed experiments, performed experiments, and analyzed data. V.S.B. and Y.J.B performed computational simulations. M.E.H., K.W.M, P.A. and W.S.Y. designed experiments, analyzed and interpreted data and edited the manuscript. V.V.L., H.T.W., V.S.B., P.M.A., Y.J.B and B.R.U wrote the manuscript. Y.J.B. and B.R.U. conceived and led the project.

## References

Akoh CC, Lee G-C, Liaw Y-C, Huang T-H, Shaw J-F (2004) GDSL family of serine esterases/lipases. Progress in Lipid Research 43 (6):534–552. doi: http://dx.doi.org/10.1016/j.plipres.2004.09.002

Anandakrishnan R, Aguilar B, Onufriev AV (2012) H++3.0: automating pK prediction and the preparation of biomolecular structures for atomistic molecular modeling and simulations. Nucleic Acids Research 40 (W1):W537–W541. doi: 10.1093/nar/gks375

Bharadwaj VS, Dean AM, Maupin CM (2013) Insights into the Glycyl Radical Enzyme Active Site of Benzylsuccinate Synthase: A Computational Study. Journal of the American Chemical Society 135 (33):12279–12288. doi: 10.1021/ja404842r

Biely P (2012) Microbial carbohydrate esterases deacetylating plant polysaccharides. Biotechnology Advances 30 (6):1575–1588. doi: http://dx.doi.org/10.1016/j.biotechadv.2012.04.010

Biely P, Mastihubová M, La Grange DC, Van Zyl WH, Prior BA (2004) Enzyme-coupled assay of acetylxylan esterases on monoacetylated 4-nitrophenyl β-d-xylopyranosides. Analytical biochemistry 332 (1):109–115

Brooks BR, Brooks CL, MacKerell AD, Nilsson L, Petrella RJ, Roux B, Won Y, Archontis G, Bartels C, Boresch S (2009) CHARMM: the biomolecular simulation program. Journal of computational chemistry 30 (10):1545–1614

Brott AS, Sychantha D, Clarke AJ (2019) Assays for the Enzymes Catalyzing the O-Acetylation of Bacterial Cell Wall Polysaccharides. In: Bacterial Polysaccharides. Springer, pp 115–136

Buller AR, Townsend CA (2013) Intrinsic evolutionary constraints on protease structure, enzyme acylation, and the identity of the catalytic triad. P Natl Acad Sci USA 110 (8):E653–E661. doi: 10.1073/pnas.1221050110

Busse-Wicher M, Gomes TC, Tryfona T, Nikolovski N, Stott K, Grantham NJ, Bolam DN, Skaf MS, Dupree P (2014) The pattern of xylan acetylation suggests xylan may interact with cellulose microfibrils as a twofold helical screw in the secondary plant cell wall of *Arabidopsis thaliana*. Plant J 79 (3):492–506. doi: 10.1111/tpj.12575

Chen VB, Arendall WB, 3rd, Headd JJ, Keedy DA, Immormino RM, Kapral GJ, Murray LW, Richardson JS, Richardson DC (2010) MolProbity: all-atom structure validation for macromolecular crystallography. Acta Crystallogr D Biol Crystallogr 66 (Pt 1):12–21. doi:S0907444909042073 [pii] 10.1107/S0907444909042073

Chong S-L, Virkki L, Maaheimo H, Juvonen M, Derba-Maceluch M, Koutaniemi S, Roach M, Sundberg B, Tuomainen P, Mellerowicz EJ (2014) O-Acetylation of glucuronoxylan in Arabidopsis thaliana wild type and its change in xylan biosynthesis mutants. Glycobiology 24 (6):494–506

Culbertson AT, Ehrlich JJ, Choe J-Y, Honzatko RB, Zabotina OA (2018) Structure of xyloglucan xylosyltransferase 1 reveals simple steric rules that define biological patterns of xyloglucan polymers. Proceedings of the National Academy of Sciences 115 (23):6064–6069

Darden T, York D, Pedersen L (1993) Particle mesh Ewald: An N· log (N) method for Ewald sums in large systems. The Journal of chemical physics 98 (12):10089–10092

Dodson G, Wlodawer A (1998) Catalytic triads and their relatives. Trends in biochemical sciences 23 (9):347–352

Emsley P, Lohkamp B, Scott WG, Cowtan K (2010) Features and development of Coot. Acta Crystallogr D Biol Crystallogr 66 (Pt 4):486–501. doi:S0907444910007493 [pii] 10.1107/S0907444910007493

Engh RA, Huber R (1991) Accurate Bond and Angle Parameters for X-Ray Protein-Structure Refinement. Acta Crystallographica Section A 47:392–400

Grantham NJ, Wurman-Rodrich J, Terrett OM, Lyczakowski JJ, Stott K, Iuga D, Simmons TJ, Durand-Tardif M, Brown SP, Dupree R (2017) An even pattern of xylan substitution is critical for interaction with cellulose in plant cell walls. Nature plants 3 (11):859

Guvench O, Hatcher E, Venable RM, Pastor RW, MacKerell AD (2009) CHARMM Additive All-Atom Force Field for Glycosidic Linkages between Hexopyranoses. Journal of Chemical Theory and Computation 5 (9):2353–2370. doi: 10.1021/ct900242e

Hendrickson WA, Teeter MM (1981) Structure of the hydrophobic protein crambin determined directly from the anomalous scattering of sulphur. Nature 290 (5802):107–113. doi: 10.1038/290107a0

Huang J, MacKerell AD (2013) CHARMM36 all-atom additive protein force field: validation based on comparison to NMR data. Journal of Computational Chemistry 34 (25):2135–2145. doi: 10.1002/jcc.23354

Humphrey W, Dalke A, Schulten K (1996) VMD: Visual molecular dynamics. Journal of Molecular Graphics 14 (1):33–38. doi: 10.1016/0263-7855(96)00018-5

Hynninen AP, Crowley MF (2014) New faster CHARMM molecular dynamics engine. Journal of computational chemistry 35 (5):406–413

Johnson AM, Kim H, Ralph J, Mansfield SD (2017) Natural acetylation impacts carbohydrate recovery during deconstruction of Populus trichocarpa wood. Biotechnology for biofuels 10 (1):48

Jorgensen WL, Chandrasekhar J, Madura JD, Impey RW, Klein ML (1983) Comparison of simple potential functions for simulating liquid water. The Journal of chemical physics 79 (2):926–935

Kabel MA, de Waard P, Schols HA, Voragen AG (2003) Location of O-acetyl substituents in xylo-oligosaccharides obtained from hydrothermally treated Eucalyptus wood. Carbohydrate research 338 (1):69–77

Kang X, Kirui A, Widanage MCD, Mentink-Vigier F, Cosgrove DJ, Wang T (2019) Lignin-polysaccharide interactions in plant secondary cell walls revealed by solid-state NMR. Nature communications 10 (1):347

Köhnke T, Östlund Å, Brelid H (2011) Adsorption of Arabinoxylan on Cellulosic Surfaces: Influence of Degree of Substitution and Substitution Pattern on Adsorption Characteristics. Biomacromolecules 12 (7):2633–2641. doi: 10.1021/bm200437m

Lassfolk R, Rahkila J, Johansson MP, Ekholm FS, Wärnå J, Leino R (2018) Acetyl group migration across the saccharide units in oligomannoside model compound. Journal of the American Chemical Society 141 (4):1646–1654

Lefebvre V, Fortabat M-N, Ducamp A, North HM, Maia-Grondard A, Trouverie J, Boursiac Y, Mouille G, Durand-Tardif M (2011) ESKIMO1 disruption in Arabidopsis alters vascular tissue and impairs water transport. PLoS One 6 (2):e16645

Ma J, Lu Q, Yuan Y, Ge H, Li K, Zhao W, Gao Y, Niu L, Teng M (2011) Crystal structure of isoamyl acetate-hydrolyzing esterase from Saccharomyces cerevisiae reveals a novel active site architecture and the basis of substrate specificity. Proteins: Structure, Function, and Bioinformatics 79 (2):662–668. doi: 10.1002/prot.22865

Manabe Y, Verhertbruggen Y, Gille S, Harholt J, Chong S-L, Pawar PM, Mellerowicz E, Tenkanen M, Cheng K, Pauly M (2013) RWA proteins play vital and distinct roles in cell wall O-acetylation in Arabidopsis thaliana. Plant physiology:pp. 113.225193

Moremen KW, Ramiah A, Stuart M, Steel J, Meng L, Forouhar F, Moniz HA, Gahlay G, Gao Z, Chapla D (2018) Expression system for structural and functional studies of human glycosylation enzymes. Nature chemical biology 14 (2):156

Morris GM, Huey R, Lindstrom W, Sanner MF, Belew RK, Goodsell DS, Olson AJ (2009) AutoDock4 and AutoDockTools4: Automated Docking with Selective Receptor Flexibility. Journal of Computational Chemistry 30 (16):2785–2791. doi: 10.1002/jcc.21256

Moynihan PJ, Clarke AJ (2013) Assay for peptidoglycan O-acetyltransferase: a potential new antibacterial target. Analytical biochemistry 439 (2):73–79

Moynihan PJ, Clarke AJ (2014) Mechanism of action of peptidoglycan O-acetyltransferase B involves a Ser-His-Asp catalytic triad. Biochemistry 53 (39):6243–6251

Murshudov GN, Skubak P, Lebedev AA, Pannu NS, Steiner RA, Nicholls RA, Winn MD, Long F, Vagin AA (2011) REFMAC5 for the refinement of macromolecular crystal structures. Acta Crystallogr D Biol Crystallogr 67 (Pt 4):355–367. doi: S0907444911001314 [pii] 10.1107/S0907444911001314

Pauly M, Ramírez V (2018) New Insights Into Wall Polysaccharide O-Acetylation. Frontiers in plant science 9

Roe DR, Cheatham TE (2013) PTRAJ and CPPTRAJ: Software for Processing and Analysis of Molecular Dynamics Trajectory Data. Journal of Chemical Theory and Computation 9 (7):3084–3095. doi: 10.1021/ct400341p

Ryckaert J-P, Ciccotti G, Berendsen HJC (1977) Numerical integration of the cartesian equations of motion of a system with constraints: molecular dynamics of n-alkanes. Journal of Computational Physics 23 (3):327–341. doi: http://dx.doi.org/10.1016/0021-9991(77)90098-5

Schultink A, Naylor D, Dama M, Pauly M (2015) The role of the plant-specific AXY9 protein in Arabidopsis cell wall polysaccharide O-acetylation. Plant physiology:pp. 114.256479

Skubak P, Pannu NS (2013) Automatic protein structure solution from weak X-ray data. Nature Communications 4. doi: Unsp 2777 Doi 10.1038/Ncomms3777

Smith PJ, Wang H-T, York WS, Peña MJ, Urbanowicz BR (2017) Designer biomass for next-generation biorefineries: leveraging recent insights into xylan structure and biosynthesis. Biotechnology for biofuels 10 (1):286

Stranne M, Ren Y, Fimognari L, Birdseye D, Yan J, Bardor M, Mollet JC, Komatsu T, Kikuchi J, Scheller HV (2018) TBL 10 is required for O-acetylation of pectic rhamnogalacturonan-I in Arabidopsis thaliana. The Plant Journal 96 (4):772–785

Sychantha D, Little DJ, Chapman RN, Boons G-J, Robinson H, Howell PL, Clarke AJ (2018) PatB1 is an O-acetyltransferase that decorates secondary cell wall polysaccharides. Nature chemical biology 14 (1):79

Urbanowicz BR, Bharadwaj VS, Alahuhta M, Peña MJ, Lunin VV, Bomble YJ, Wang S, Yang JY, Tuomivaara ST, Himmel ME, Moremen KW, York WS, Crowley MF (2017) Structural, mutagenic and in silico studies of xyloglucan fucosylation in Arabidopsis thaliana suggest a water-mediated mechanism. The Plant Journal 91 (6):931–949. doi: doi:10.1111/tpj.13628

Urbanowicz BR, Peña MJ, Moniz HA, Moremen KW, York WS (2014) Two Arabidopsis proteins synthesize acetylated xylan in vitro. The Plant Journal 80 (2):197–206. doi: 10.1111/tpj.12643

Vanommeslaeghe K, Hatcher E, Acharya C, Kundu S, Zhong S, Shim J, Darian E, Guvench O, Lopes P, Vorobyov I, MacKerell AD (2010) CHARMM General Force Field (CGenFF): A force field for drug-like molecules compatible with the CHARMM all-atom additive biological force fields. Journal of computational chemistry 31 (4):671–690. doi: 10.1002/jcc.21367

Winn MD, Ballard CC, Cowtan KD, Dodson EJ, Emsley P, Evans PR, Keegan RM, Krissinel EB, Leslie AG, McCoy A, McNicholas SJ, Murshudov GN, Pannu NS, Potterton EA, Powell HR, Read RJ, Vagin A, Wilson KS (2011) Overview of the CCP4 suite and current developments. Acta Crystallogr D Biol Crystallogr 67 (Pt 4):235–242. doi:S0907444910045749 [pii] 10.1107/S0907444910045749

Xiong G, Cheng K, Pauly M (2013) Xylan O-acetylation impacts xylem development and enzymatic recalcitrance as indicated by the Arabidopsis mutant *tbl29*. Mol Plant 6 (4):1373–1375. doi: 10.1093/mp/sst014

Yuan Y, Teng Q, Zhong R, Haghighat M, Richardson EA, Ye Z-H (2016a) Mutations of Arabidopsis TBL32 and TBL33 affect xylan acetylation and secondary wall deposition. PLoS One 11 (1):e0146460

Yuan Y, Teng Q, Zhong R, Ye Z-H (2015) TBL3 and TBL31, two Arabidopsis DUF231 domain proteins, are required for 3-O-monoacetylation of xylan. Plant and Cell Physiology 57 (1):35–45

Yuan Y, Teng Q, Zhong R, Ye ZH (2016b) Roles of Arabidopsis TBL34 and TBL35 in xylan acetylation and plant growth. Plant Sci 243:120–130. doi: 10.1016/j.plantsci.2015.12.007

Zhong R, Cui D, Richardson EA, Phillips DR, Azadi P, Lu G, Ye Z-H (2019) Cytosolic Acetyl-CoA Generated by ATP-Citrate Lyase Is Essential for Acetylation of Cell Wall Polysaccharides. Plant and Cell Physiology

Zhong R, Cui D, Ye Z-H (2017) Regiospecific acetylation of xylan is mediated by a group of DUF231-containing O-acetyltransferases. Plant and Cell Physiology 58 (12):2126–2138

Zhong R, Cui D, Ye Z-H (2018) Xyloglucan O-acetyltransferases from Arabidopsis thaliana and Populus trichocarpa catalyze acetylation of fucosylated galactose residues on xyloglucan side chains. Planta 248 (5):1159–1171

